# A nuclear targeting approach enables efficient transgenesis in the milkweed bug *Oncopeltus fasciatus*

**DOI:** 10.64898/2026.07.30.741915

**Authors:** Yu Shirai, Yousuf Hashmi, Ashley Watts, Jonchee A. Kao, Cassandra G. Extavour

## Abstract

Insects show extreme diversity and have long intrigued biologists. Although gene editing and transgenesis should enable the use of almost any insect species for biological research, species-specific challenges remain and conditions must be optimized carefully. Here, using the milkweed bug *Oncopeltus fasciatus*, we first generate a useful eye- and body-color mutant strain by CRISPR/Cas9-mediated genome editing. Using this strain, we develop an efficient *piggyBac*-mediated transgenesis system with a nuclear targeting approach. Incorporating 1× and 3× nuclear localization signals into *piggyBac* mRNA substantially enhances overall transformation efficiency in *O. fasciatus*. Taking advantage of the mutant strain and the efficient transgenesis system, we integrated multiple expression cassettes ranging from 1.8 to 7.2 kb, including attP strains for phiC31-mediated integration and histone-labelled strains for live imaging. We further show that the Q system, a binary expression system, is functional in this species, paving the way for sophisticated genetic manipulations. Together, our results expand the genetic toolkit of *O. fasciatus* as a comparative model insect and provide a practical framework for developing efficient transgenesis in other non-traditional model organisms.

**Summary statement:** We developed an efficient *piggyBac* transgenesis system in the milkweed bug with a nuclear targeting approach, expanding genetic tools for non-model insects.

## Introduction

Insects are one of the most diverse groups of animals on Earth (Stork, 2017). Their remarkable beauty, adaptations, and diversity have long fascinated biologists. Many biological questions about insect diversity and evolution cannot be addressed using the long-standing arthropod model *Drosophila melanogaster* alone (Clark et al., 2019; Kao et al., 2026; Truman et al., 2026). Beyond the *Drosophila melanogaster* paradigm, recent technological advancements have enabled us to use functional genetics to investigate many non-traditional model organisms, opening new avenues for studies of the evolution of developmental mechanisms (Gudmunds et al., 2022). Comparative studies are essential to understand the evolutionary history of insects. Establishing genomic and biological resources across multiple species has thus become a critical investment in cell and developmental biology.

The large milkweed bug, *Oncopeltus fasciatus* (Lygaeidae, Hemiptera) has a long history of comparative developmental studies (reviewed in Chipman, 2017; Feir, 1974). Hemiptera is a large and diverse order of insects that is phylogenetically important for understanding insect evolution, as it is an outgroup to Holometabola, the clade of insects that undergo complete metamorphosis (Misof et al., 2014). In the mid-twentieth century, *O. fasciatus* was widely used as a model organism across many different contexts, including embryology, physiology, and histology (e.g. Butt, 1949; Kelly and Telfer, 1979; Schreiner, 1977). However, the rise of *D. melanogaster* genetics largely displaced *O. fasciatus*, along with other insect species. Approximately two decades ago, taking advantage of its ease of rearing and high sensitivity to RNAi-mediated gene perturbation, *O. fasciatus* began to re-emerge as an evo-devo model, particularly for the study of segmentation and body patterning (Angelini and Kaufman, 2005; Angelini et al., 2005; Liu and Kaufman, 2004a; Liu and Kaufman, 2004b; Liu and Kaufman, 2005; Liu and Patel, 2009; Liu and Patel, 2010). More recently, the development of CRISPR/Cas9-mediated targeted mutagenesis (Reding and Pick, 2020) and knock-in methods (Shirai et al., 2026) has further expanded its utility as a comparative model organism for functional genetics.

Since transgenesis is a powerful genetic tool for investigating gene function and *cis*-regulatory elements (Gregory et al., 2016; Handler, 2002), deploying this tool in *O. fasciatus* is essential for establishing it as a more versatile model organism. We focus here on a hyperactive variant of the *piggyBac* transposase called hyPBase (Yusa et al., 2011), which has been successfully applied across multiple insect species (Chen and Palli, 2021; Gonzalez-Sqalli et al., 2024; Hart et al., 2023; Heryanto et al., 2022; Inada et al., 2025; Kou et al., 2023; Ohde et al., 2025), consistently outperforming the original *piggyBac* transposase in overall efficiency (Eckermann et al., 2018; Ohde et al., 2025; Otte et al., 2018).

Here, we describe an efficient *piggyBac*-mediated transgenesis system in *O. fasciatus*. First, we generate a useful eye- and body-color mutant strain by crossing the *Of-vermilion* (*Of-v*) mutant (Reding et al., 2023) with a newly generated CRISPant strain carrying a mutation in *Of-xanthine dehydrogenase* (*Of*-*xdh1*), the ortholog of *D. melanogaster rosy*, enabling easier detection of fluorescent signals through the adult cuticle. Using this double mutant strain, we develop a highly efficient *piggyBac*-mediated transgenesis system by incorporating additional SV40 nuclear localization signals (NLSs) (Kalderon et al., 1984) into hyPBase. Furthermore, we provide evidence that Q system mediated-binary expression (Potter et al., 2010) is functional in *O. fasciatus*. Together, our results substantially expand the experimental possibilities of *O. fasciatus* as a comparative genetic model organism and provide a practical framework for implementing efficient transgenesis in other non-traditional model organisms.

## Results

### Establishment of a useful eye- and body-color mutant strain in *O. fasciatus*

Many successful insect transgenesis systems rely on detection of a dominant marker, often expression of a fluorescent protein, in the eye (Berghammer et al., 1999). However, detecting this marker expression can be challenging if the animal’s eyes are darkly pigmented. In the case of *O. fasciatus*, the wild-type eye color is black. Therefore, we aimed to generate a mutant strain with a lighter eye color to facilitate the detection of fluorescent signal, such as GFP or RFP, derived from transgenesis. Previous studies have attempted to generate such a strain: one study reported that the *O. fasciatus white* (*Of-w*) gene, commonly used to provide a suitable background for detection of a dominant rescue construct in other insects, caused lethality when disrupted (Reding and Pick, 2020), while another study established strains mutant for *Of-v*, an ommochrome pathway gene, displaying dark red eye color (Reding et al., 2023). However, we were not certain whether the eyes of the *Of*-*v* mutants were light enough to detect fluorescent signal. In 1970, Peter Lawrence reported *O. fasciatus* mutants named "*red eye* (*re*)", "*white body* (*wb*)", and "*cream body* (*cb*)" (Lawrence, 1970). *re* mutants had red eyes, and the described phenotypes of *re* mutants are generally consistent with that of *Of-v* mutants (Reding et al., 2023), suggesting that *re* mutants lack ommochrome pigments. However, *re* was reported to be autosomal (Lawrence, 1970), whereas *Of-v* is located on the X-chromosome (Reding et al., 2023), indicating that the gene responsible the *re* phenotype is different from *Of-v*. By contrast, *wb* and *cb* mutants were both reported to have a whitish yellow body color, and these described phenotypes are comparable to the loss of orange body coloration observed upon systemic RNAi of pteridine pathway genes (Liu, 2016; Reding et al., 2023). Importantly, double mutants of *re* and either *wb* or *cb* showed eye color distinct from that of the single mutants (Lawrence, 1970), and maternal RNAi of *punch*, a pteridine pathway gene, resulted in eye-color defects in embryos (Liu, 2016). Together, these results suggested that both ommochrome and pteridine pigments might contribute to eye color formation in *O. fasciatus*. Based on these findings, we decided to target the *O. fasciatus* ortholog of the pteridine pathway gene *xdh1* (*Of-xdh1*, Accession No. OFAS027123), whose *D. melanogaster* ortholog is the *rosy* gene (Keith et al., 1987). *Of-xdh1* has been implicated in body coloration in *O. fasciatus* based on systemic RNAi-mediated functional analysis (Reding et al., 2023).

*Of-xdh1* is predicted to encode a 1341 amino acid pteridine pathway enzyme containing 2Fe-2S, flavin adenine dinucleotide (FAD) binding, and molybdopterin binding domains, as determined by InterProScan (Fig. 1A). To generate a homozygous mutant *Of-xdh1* strain (Fig. 1B), we designed a guide RNA targeting the region immediately downstream of the start codon (Fig. 1A), hoping to generate an indel and frameshift resulting in a premature stop codon and eliminating the functional domains of this enzyme. Following injection of Cas9/gRNA ribonucleoprotein complex into 54 eggs, 17 of 18 hatchlings (94.4%) showed a clear body color difference in the injected generation (G_0_) compared to wild type (WT), enabling selection of potential mutants (Fig. 1C). After three rounds of crossing, we established a homozygous strain named *xdh1*^Δ*2*^, carrying a two base pair deletion allele introducing a premature stop codon, and displaying orange body coloration (Fig. 1D–E and Supplementary Fig. 1).

**Figure 1.**
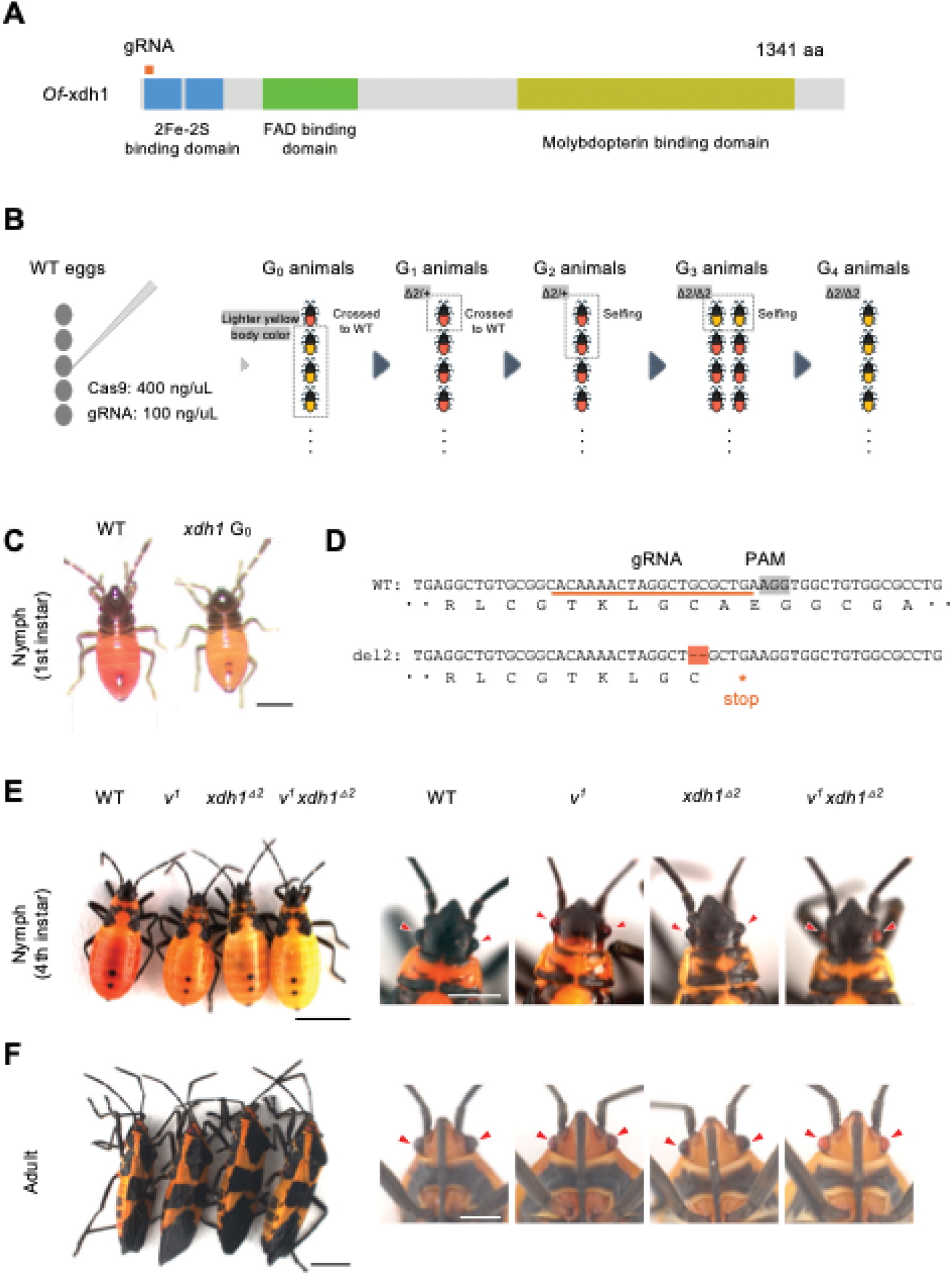
Establishment of a useful double mutant strain in *O. fasciatus*. (A) *Of*-xdh1 predicted protein structure. The N-terminal orange square indicates the gRNA target site. Protein domains were predicted using InterProScan. FAD, flavin adenine dinucleotide. (B) Schematic of the experimental workflow and genetic crosses performed to establish the homozygous *xdh1* mutant strain. (C) Representative images of wild type (WT) and *xdh1* G_0_ mutant hatchlings (dorsal view). Scale bar, 500 μm. (D) gRNA target site and mutant allele of *xdh1*. (E) Representative images of nymphs (dorsal view) and close-up views of eye coloration (dorsal view) in WT, *v^1^*, *xdh1*^Δ*2*^, and *v^1^ xdh1*^Δ*2*^. Arrowheads indicate eye color. Black scale bar, 3 mm; white scale bar, 1 mm. (F) Representative images of adults (lateral view) and close-up views of eye coloration (ventral view) in WT, *v^1^*, *xdh1*^Δ*2*^, and *v^1^ xdh1*^Δ*2*^. Arrowheads indicate eye color. Black scale bar, 3 mm; white scale bar, 1 mm. Anterior is up in (C), (E), and (F).

To investigate the potential involvement of *xdh1* in eye-color formation, we crossed *xdh1*^Δ*2*^ animals with those from the *Of-v* mutant strain *v^1^* (Reding et al., 2023) (Supplementary Fig. 2) to generate a double homozygous mutant strain. We confirmed the mutations at both loci by sequencing. Comparison of eye color among four different strains (wild type, *v^1^*, *xdh1*^Δ*2*^, and *v^1^ xdh1*^Δ*2*^), revealed that the *v^1^ xdh1*^Δ*2*^ double mutant strain displayed a brighter orange eye color than all the others (Fig. 1E–F and Supplementary Fig. 3). The double mutant strain also showed a brighter yellowish body color than the others, particularly at early nymphal stages (Fig. 1E, and Supplementary Fig. 3), indicating that both ommochrome and pteridine pigments contribute to eye and body color formation in *O. fasciatus*. Thus, we used this strain for the subsequent transgenesis experiments described in the following sections.

### Nuclear targeting allows an efficient hyPBase-mediated transgenesis in *O. fasciatus*

To develop a transgenesis method in *O. fasciatus* using the *v^1^ xdh1*^Δ*2*^ strain (Fig. 2A), we first utilized the *piggyBac* vector *pXL[9×P3-mScarlet-I-SV40T]* (1,775 bp). In this vector, mScarlet-I is placed under the control of the 9×P3 promoter, consisting of nine copies of a Pax6 homeodomain binding site, which drives specific expression in the eyes in many insects (Berghammer et al., 1999; Horn and Wimmer, 2000; Horn et al., 2000; Inada et al., 2025; Ohde et al., 2025; Sheng et al., 1997) (Fig. 2B). We screened for mScarlet-I expression in the eyes of injected animals at both mid-nymphal and adult stages. In the first round of injection with *in vitro* transcribed *hyPBase* mRNA, 5/29 G_0_ animals were mScarlet-I-positive at nymphal stages (G_0_ somatic transformation rate (STR) = 17.2%), and 6/25 (STR = 24.0%) at adult stages (Fig. 2C–D and Table 1). However, no G_0_ animals showed mScarlet-I expression at either stage in a second round of injection using the same batch of reagents (0/70 at nymphal stages and 0/62 at adult stages; STR = 0%) (Fig. 2D and Table 1), suggesting inconsistent transgenesis efficiency with unmodified hyPBase.

**Figure 2.**
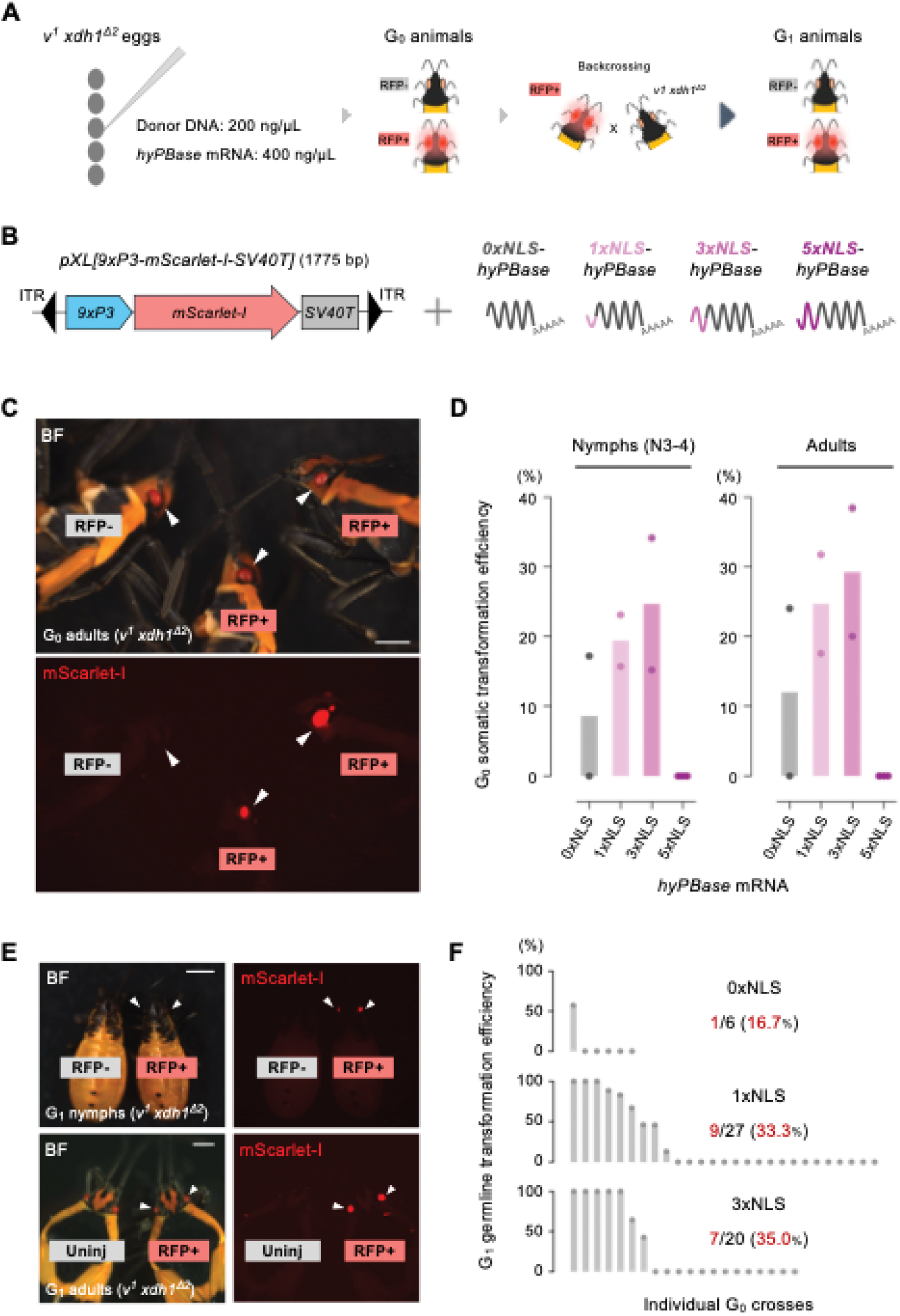
Investigation of a nuclear targeting approach for hyPBase-mediated transgenesis in *O. fasciatus*. (A) Schematic of the experiments to identify G_0_ mScarlet-I-positive animals and establish stable transgenic lines. (B) Schematic of the *piggyBac* donor vector *pXL[9×P3-mScarlet-I-SV40T]* and *hyPBase* mRNA carrying 0×, 1×, 3×, or 5×NLS. (C) Representative images of G_0_ mScarlet-I-positive adults. Scale bar, 1 mm. (D) Mean G_0_ somatic transformation rate (STR) across *hyPBase* mRNA carrying 0×, 1×, 3×, or 5×NLS. Each dot represents an independent injection experiment. (E) Representative images of G_1_ mScarlet-I-positive nymphs and adults. Arrowheads indicate eyes. Scale bar, 1 mm. (F) Mean G_1_ germ line transformation efficiency (GTE). The height of each bar represents the mean proportion of mScarlet-I-positive animals among G_1_ progeny from an individual cross. Each dot represents an individual backcross that yielded G_1_ progeny. Red numbers indicate the number of crosses yielding heritable mutant alleles and the corresponding percentages.

**Table 1.** Detailed results of *piggyBac*-mediated transgenesis experiments with *hyPBase* mRNA carrying 0×, 1×, 3×, or 5×NLS in *O. fasciatus*. Each row shows the results of an independent experiment. All G_0_ nymphs and adults reached each respective stage were screened. Percentages shown in parentheses indicate somatic transformation rates (STR). Nymphs were screened at stages 3 to 5.

| **Donor plasmid** | **Transposase (mRNA)** | **Injected eggs** | **% injected eggs survived to nymphal screening** | **Screened G0 nymphs** | **mScarlet-I + nymphs** | **mScarlet-I - nymphs** | **% screened nymphs survived to adult screening** | **Screened G0 adults** | **mScarlet-I + adults** | **mScarlet-I - adults** |
|---|---|---|---|---|---|---|---|---|---|---|
| pXL[9xP3-mScarlet-I-SV40T] | 0xNLS-hyPBase | 135 | 21.5% | 29 | 5 (17.2%) | 24 | 86.2% | 25 | 6 (24.0%) | 19 |
| pXL[9xP3-mScarlet-I-SV40T] | 0xNLS-hyPBase | 246 | 28.5% | 70 | 0 | 70 | 88.6% | 62 | 0 | 62 |
| pXL[9xP3-mScarlet-I-SV40T] | 1xNLS-hyPBase | 170 | 45.9% | 78 | 18 (23.1%) | 60 | 80.8% | 63 | 20 (31.8%) | 43 |
| pXL[9xP3-mScarlet-I-SV40T] | 1xNLS-hyPBase | 185 | 37.8% | 70 | 11 (15.7%) | 59 | 81.4% | 57 | 10 (17.5%) | 47 |
| pXL[9xP3-mScarlet-I-SV40T] | 3xNLS-hyPBase | 161 | 28.6% | 46 | 7 (15.2%) | 39 | 89.1% | 41 | 8 (20.0%) | 33 |
| pXL[9xP3-mScarlet-I-SV40T] | 3xNLS-hyPBase | 218 | 20.1% | 44 | 15 (34.1%) | 29 | 90.8% | 40 | 15 (38.5%) | 25 |
| pXL[9xP3-mScarlet-I-SV40T] | 5xNLS-hyPBase | 130 | 34.6% | 45 | 0 | 45 | 86.7% | 39 | 0 | 39 |
| pXL[9xP3-mScarlet-I-SV40T] | 5xNLS-hyPBase | 80 | 7.5% | 6 | 0 | 6 | 83.3% | 5 | 0 | 5 |
| pXL[9xP3-mScarlet-I-SV40T] | 5xNLS-hyPBase | 73 | 23.3% | 17 | 0 | 17 | 94.1% | 16 | 0 | 16 |

Given this inconsistency across technical replicate experiments, we hypothesized that incorporating additional SV40T nuclear localization signals (NLSs) into hyPBase would improve transgenesis efficiency (Fig. 2B). To test this hypothesis, we compared STR values across 0×, 1×, 3× and 5×NLS *hyPBase* mRNA conditions using the same donor vector. We found that 1× and 3×NLS *hyPBase* mRNA yielded higher and more consistent STR values than 0×NLS. Across two experimental replicates, STR values for 1×NLS ranged from 15.7% (11/70) to 23.1% (18/78) at nymphal stages and from 17.5% (10/57) to 31.8% (20/63) at adult stages; 3×NLS ranged from 15.2% (7/46) to 34.1% (15/44) at nymphal stages and from 20.0% (8/41) to 38.5% (15/40) at adult stages (Fig. 2D and Table 1). By contrast, with 5×NLS we recovered no positive G_0_ animals (0/68 at nymphal stages) (Fig. 2D and Table 1). We also noted that screening at nymphal stages underestimates STR values (Fig. 2D and Table 1), possibly because the eye color of the *v^1^ xdh1*^Δ*2*^ strain becomes progressively lighter with development (Fig. 1E–F and Supplementary Fig. 3).

To investigate whether this trend was conserved in another insect species, we injected the same series of *hyPBase* mRNAs with the same donor vector *pXL[9xP3-mScarlet-I-SV40T]* into embryos of the cricket *Gryllus bimaculatus* (Supplementary Fig. 4 and Supplementary Table 1). In contrast to our results from *O. fasciatus* (Fig. 2D and Table 1), we found no differences in efficiency across NLS copy number in *G. bimaculatus* (Supplementary Fig. 4 and Supplementary Table 1), although overall efficiency in *G. bimaculatus* was generally higher than in *O. fasciatus* (Table 1 and Supplementary Table 1). Together, these results indicate that incorporating 1× and 3×NLS into *hyPBase* mRNA may be a practical strategy to improve transgenesis efficiency in species where standard methods are inefficient.

To assess germ line transmission, we backcrossed mScarlet-I-positive G_0_ *O. fasciatus* adults with *v^1^ xdh1*^Δ*2*^ adults of the opposite sex and counted the number of mScarlet-I-positive G_1_ animals emerging from single pair matings (see Supplementary Tables 2–4 for detailed results of individual crosses). In positive G_1_ animals we detected mScarlet-I expression throughout the compound eyes and in the ocelli (Fig. 2E). G_1_ germ line transformation efficiency (GTE) with 0×NLS *hyPBase* mRNA was 16.7% (1/6 crosses), whereas 1× and 3×NLS conditions provided approximate twofold enhancement, yielding 33.3% (9/27 crosses) and 35.0% (7/20 crosses) GTE, respectively (Fig. 2F and Supplementary Tables 2–4).

To determine whether the *v^1^ xdh1*^Δ*2*^ mutant background indeed facilitates the detection of mScarlet-I expression in the eyes, we compared mScarlet-I visibility between wild type and *v^1^ xdh1*^Δ*2*^ backgrounds. We found that mScarlet-I expression was visible throughout the eye in the *v^1^ xdh1*^Δ*2*^ background, whereas in the wild-type background the visible signal was restricted to a small, dot-like area at the center of the eye (Supplementary Fig. 5). These results confirm that the *v^1^ xdh1*^Δ*2*^ background enhances marker visibility.

### Evaluation of a 9×P3-EGFP transgenesis marker in *O. fasciatus*

To expand the transgenesis toolkit in *O. fasciatus*, we next tested the donor plasmid *pXL[9×P3-EGFP-SV40T, attP]* (1,838 bp) carrying an EGFP transformation marker driven by the 9×P3 promoter and an attP site for future application of phiC31-mediated site-specific integration (Fig. 3A) (Groth et al., 2004). Two experimental replicates of injection with 3×NLS *hyPBase* mRNA resulted in 15.4% (10/65) and 9.3% (5/54) G_0_ STR when screened at nymphal stages (Table 2). As with mScarlet-I expression, we observed EGFP expression in the compound eyes and in the ocelli (Fig. 3B). Among successful backcrosses (12 of 15 EGFP-positive G_0_ animals) with animals of the opposite sex from the *v^1^ xdh1*^Δ*2*^ strain, we recovered EGFP-positive G_1_ animals from each of five independent G_0_ crosses (5/12 crosses; GTE = 41.7%) (Fig. 3C–D; see Supplementary Table 5 for detailed results of individual crosses). We then crossed four G_1_ adults with animals of the opposite sex of the *v^1^ xdh1*^Δ*2*^ strain and found that three of these crosses showed >72.5% G_2_ transgene inheritance rates, suggesting that the parent G_1_ animals were likely to have more than one transgene (Supplementary Table 5). Using two G_2_ animals from one of the crosses which showed a 50% inheritance rate, consistent with one insertion, we performed splinkerette PCR (Potter and Luo, 2010) and determined the transgene insertion site by sequencing the resulting PCR products, confirming genomic integration of the transgene (Supplementary Fig. 6).

**Figure 3.**
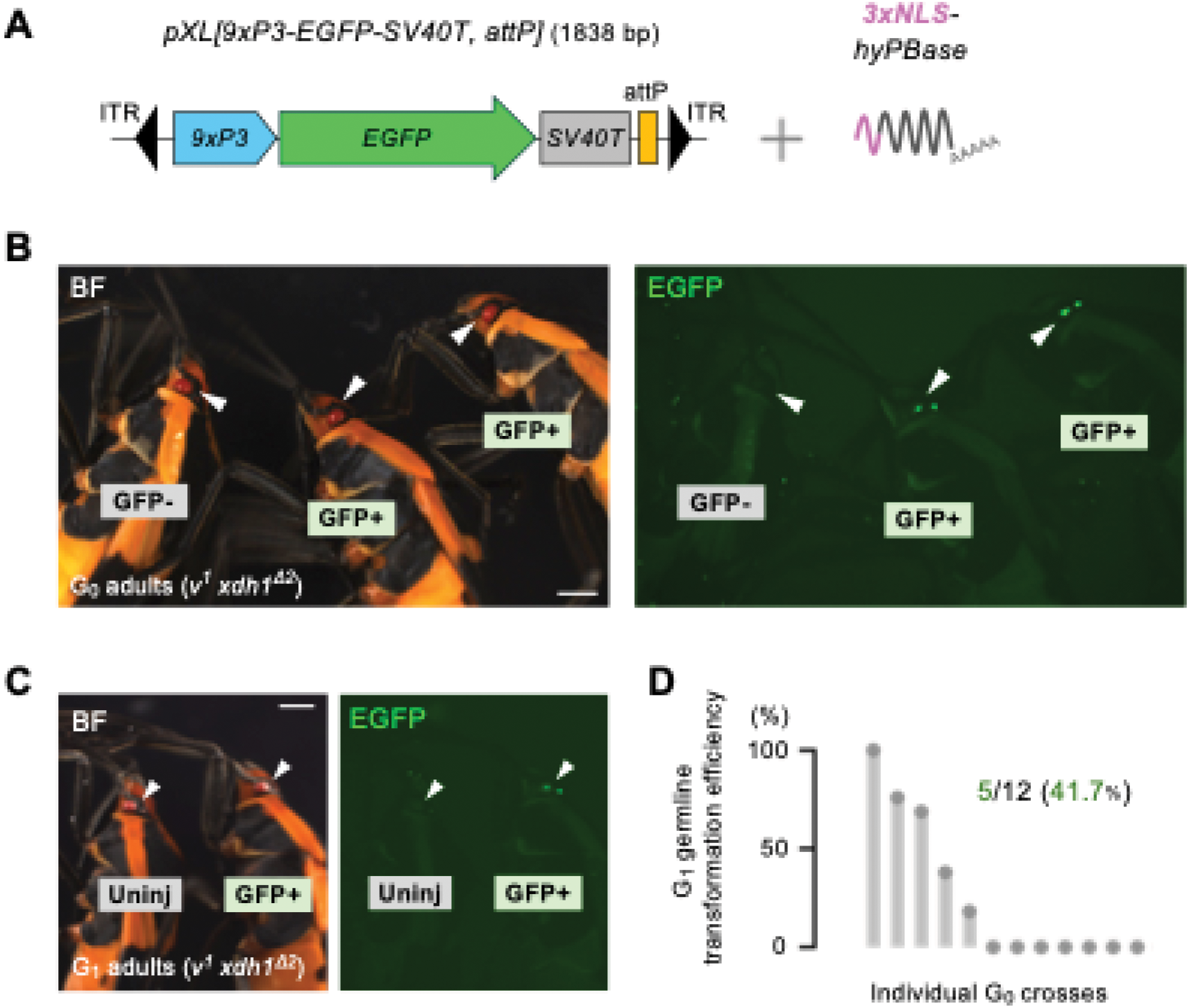
Establishment of multiple attP strains using 3×NLS *hyPBase* in *O. fasciatus.* (A) Schematic of the *piggyBac* donor vector *pXL[9×P3-EGFP-SV40T, attP]* and *hyPBase* mRNA carrying 3×NLS. (B) Representative images of G_0_ EGFP-positive adults. Arrowheads indicate eyes. Scale bar, 1 mm. (C) Representative images of G_1_ EGFP-positive adults. Arrowheads indicate eyes. Scale bar, 1 mm. (D) G_1_ germ line transformation efficiency (GTE). Each bar represents the proportion of EGFP-positive animals among G_1_ progeny from an individual cross. Each dot represents an individual backcross that yielded G_1_ progeny. Green numbers indicate the number of crosses yielding heritable transgenes and the corresponding percentages.

**Table 2.**
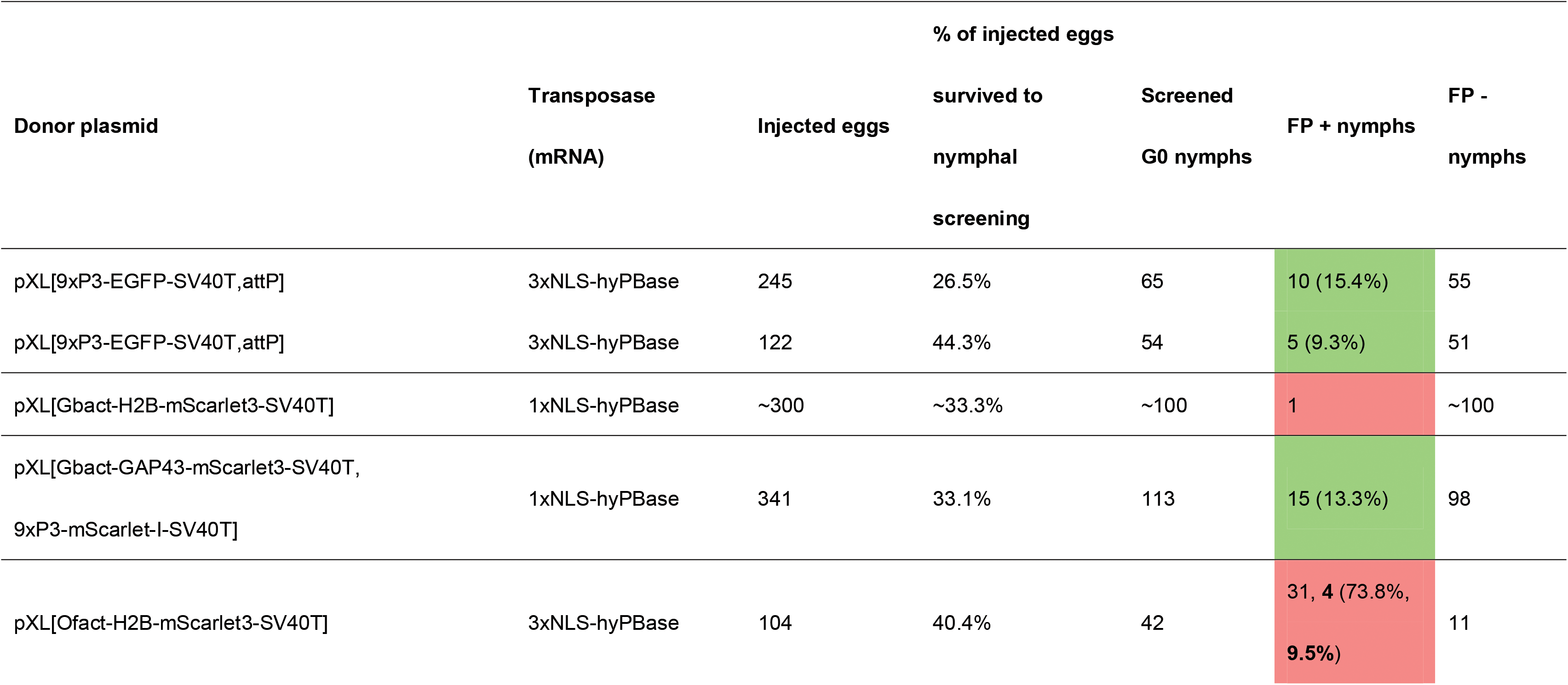
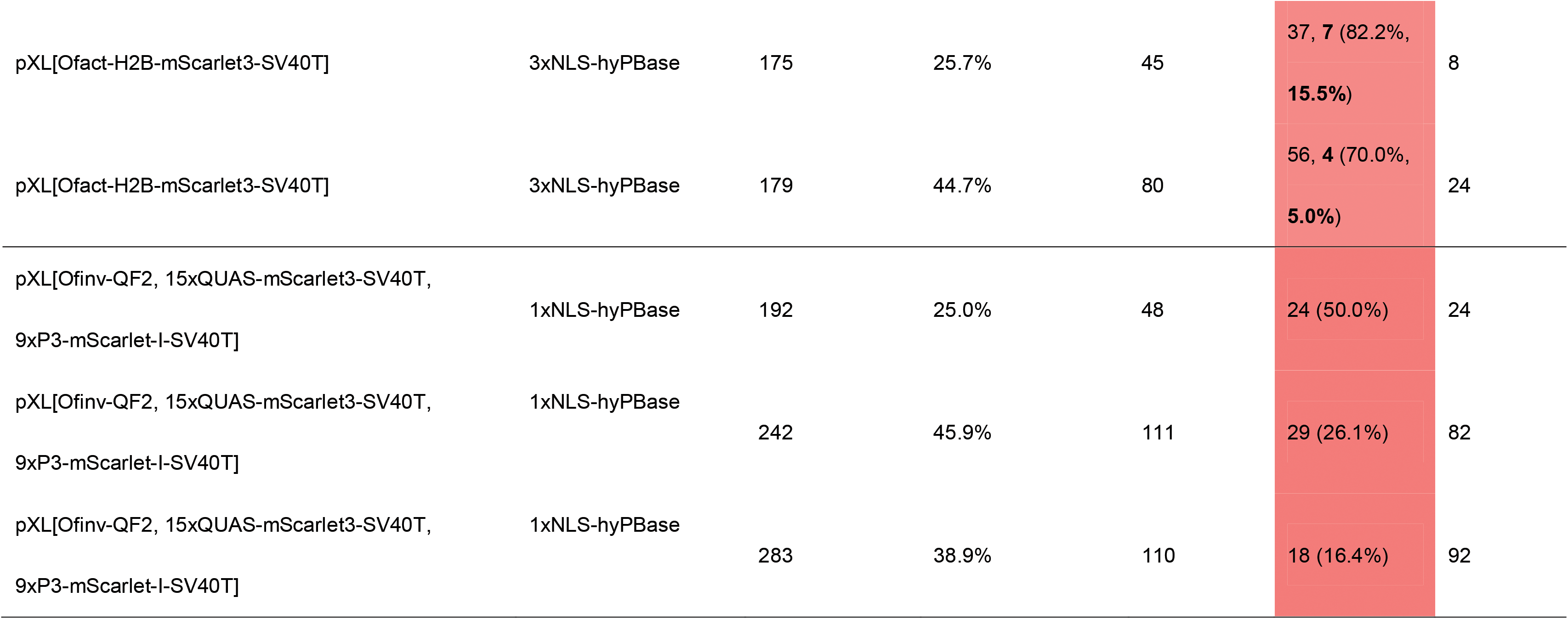
Detailed results of *piggyBac*-mediated transgenesis experiments with NLS-tagged hyPBase in *O. fasciatus*. Each row shows the results of an independent experiment. All G_0_ nymphs and adults reached each respective stage were screened. Percentages shown in parentheses indicate somatic transformation rates (STR). FP, fluorescent protein. For experiments using donor plasmid pXL[Ofact-H2B-mScarlet3-SV40T], numbers in bold indicate FP + nymphs that had detectable fluorescence in >50% of their body area. Nymphs were screened at nymphal stages 3 to 5, except for the experiment using donor plasmid pXL[Ofinv-QF2, 15xQUAS-mScarlet3-SV40T, 9xP3-mScarlet-I-SV40T], which were screened at nymphal stages 2 to 5.

### Attempt to generate a histone-labelled and a membrane-labelled transgenic milkweed bug

Nuclear and plasma membrane markers are practical and useful tools for developmental biology (e.g. Benton et al., 2013; Karapidaki et al., 2026; Nakamura et al., 2010). To try to generate a transgenic strain with ubiquitously constitutively fluorescently labelled histones, we prepared the donor plasmid *pXL[Gbact-H2B-mScarlet3-SV40T]* (2,974 bp), in which H2B-mScarlet3 is driven by the previously characterized *Gryllus bimaculatus* actin promoter (*Gbact*) (Supplementary Fig. 7). This promoter can drive ubiquitous constitutive expression and has been widely used in the cricket community (Nakamura et al., 2010; Ohde et al., 2025; Zhang et al., 2002). We reasoned that, just as some *D. melanogaster* promoters retain their function in transgenic insects of other species (Bottino-Rojas and James, 2022), the cricket promoter might retain its function in another hemimetabolous insect. We injected the donor plasmid with 1×NLS *hyPBase* mRNA and screened approximately 100 hatched G_0_ animals for mScarlet3 expression (Table 2). We obtained one animal displaying clear mScarlet3 expression through the integument across the whole body and backcrossed it with the *v^1^ xdh1*^Δ*2*^ strain to generate G_1_ progeny. We found that 42.4% of G_1_ animals (14/33) displayed the same mScarlet3 expression pattern as the G_0_ parent (Supplementary Fig. 7 and Supplementary Table 6). We observed that nuclei with detectable mScarlet3 expression were located underneath epidermal cells, whereas nuclei in overlying epidermal cells did not express detectable mScarlet3 (Supplementary Fig. 7). When we dissected ovaries from G_1_ adult females, we detected nuclear mScarlet3 expression in the ovariole muscle sheath but not in the tropharium or follicle cells (Supplementary Fig. 7). Similarly, we observed mScarlet3 expression in the testis muscle sheath, but not in sperm cells (Supplementary Fig. 7). These expression patterns suggest that the transgene may have integrated in the vicinity of a muscle-specific enhancer, resulting in enhancer trap-like expression driven by an endogenous regulatory element, rather than the ubiquitous constitutive expression driven by this *Gbact* promoter in other transgenic contexts (Nakamura et al., 2010). Importantly, this result also suggests that the *Gbact* promoter retains basal promoter activity in *O. fasciatus*, as it appears sufficient to support enhancer trap-mediated expression. We used splinkerette PCR to identify the transgene insertion site as being within an intron of an uncharacterized gene locus encoding a protein with a predicted transmembrane domain (Accession No. OFAS019046-RA; Supplementary Fig. 7).

We next aimed to generate a membrane-labelled transgenic strain using the GAP43 membrane localization tag, which has recently been shown to be broadly applicable across a wide variety of animals (Karapidaki et al., 2026). We prepared the donor plasmid *pXL[Gbact-GAP43-mScarlet3-SV40T, 9×P3-EGFP-SV40T]* (3,635 bp), in which membrane-tagged mScarlet3 and EGFP are driven by the *Gbact* and 9×P3 promoters, respectively (Supplementary Fig. 8). We injected this donor plasmid together with 1×NLS *hyPBase* mRNA and screened for EGFP expression in the eyes at nymphal stages. Of 113 screened G_0_ nymphs, 15 showed EGFP expression (STR = 13.3%) (Table 2). We backcrossed these 15 EGFP-positive G_0_ animals with animals of the opposite sex from the *v^1^ xdh1*^Δ*2*^ strain, and seven of the crosses successfully yielded G_1_ progeny. Of these, three yielded EGFP-positive G_1_ animals (GTE = 42.9%) (Supplementary Fig. 8 and Supplementary Table 7). However, we could not detect clear membrane-labelled expression of mScarlet3 in the ovary or testis of any G_1_ progeny. We did, however, detect mScarlet3 expression in the eyes of G_1_ adults (Supplementary Fig. 8), which we speculate is due to 9×P3 enhancer trapping within the transgene. Together, these results suggest that the *Gbact* promoter is active in *O. fasciatus* but may not be sufficient to drive ubiquitous expression.

### Identification of a constitutive and ubiquitous promoter in *O. fasciatus*

To identify a promoter that would drive stable, constitutive, and ubiquitous expression in *O. fasciatus*, we focused on an endogenous actin promoter. Based on a BLAST search using *D. melanogaster* Actin 5c (Act5c; NP_001284915.1), we identified two orthologs in *O. fasciatus*, *Of*-*act5ca* (Accession No. OFAS010390-RA) and *Of*-*act5cb* (Accession No. OFAS010391-RA), which are annotated as tandem loci in the genome (Panfilio et al., 2019). We prepared the donor plasmid *pXL[Ofact-H2B-mScarlet3-SV40T]* (4,650 bp), in which H2B-mScarlet3 is driven by the 2.5 kb upstream of the *Of-act5ca* gene (Fig. 4A). Three experimental replicates of injection with 3×NLS *hyPBase* mRNA resulted in 73.8% (31/42), 82.2% (37/45), and 70.0% (56/80) G_0_ STR respectively when screened at nymphal stages (Table 2). Some mScarlet3-positive G_0_ animals showed mScarlet3 expression only as dots or patches on their bodies. However, a subset of G_0_ animals exhibited mScarlet3 expression across more than 50% of their bodies from embryonic stages (Fig. 4B), which was maintained through nymphal stages. Specifically, 9.5% (4/42), 15.5% (7/45), and 5.0% (4/80) of screened nymphs displayed mScarlet3 expression across more than 50% of their bodies (Table 2). We selected nine of these strongly expressing G_0_ animals for backcrossing with animals of the opposite sex from the *v^1^ xdh1*^Δ*2*^ strain, and seven of the crosses yielded G_1_ progeny. Of these, three yielded mScarlet3-positive G_1_ animals (GTE = 42.9%) (Supplementary Table 8) displaying ubiquitous mScarlet3 expression throughout the body (a representative image at the fifth nymphal stage is shown in Fig. 4C).

**Figure 4.**
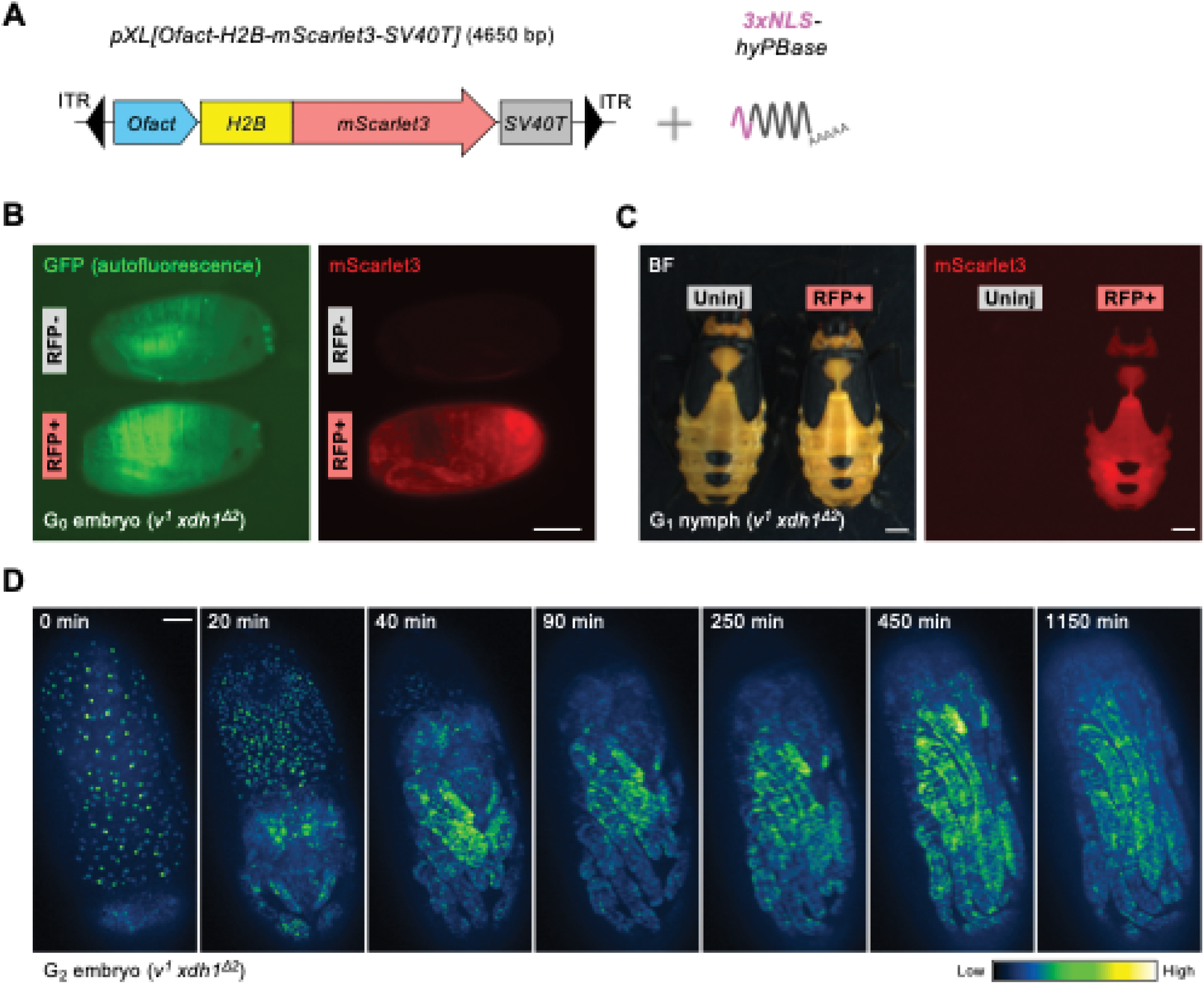
Establishment of the ubiquitous histone-labelled strain using 3×NLS *hyPBase* in *O. fasciatus*. (A) Schematic of the *piggyBac* donor vector *pXL[Ofact-H2B-mScarlet3-SV40T]* and *hyPBase* mRNA carrying 3×NLS. (B) Representative images of G_0_ mScarlet3-positive adults (lateral view). Anterior is to the right. Scale bar, 300 μm. (C) Representative images of G_1_ mScarlet3-positive nymphs (dorsal view). Anterior is up. Scale bar, 3 mm. (D) Time series of embryonic development of mScarlet3-positive G_2_ animal (ventral view), visualized with the "Green Fire Blue" relative intensity lookup table in Fiji (Schindelin et al., 2012). Maximum-intensity projections of a ventro-lateral view of the same embryo are timed relative to the first time point shown at the far left (0 min). Anterior is up. Scale bar, 100 μm.

Having successfully established a ubiquitously expressing histone-labelled transgenic *O. fasciatus*, we asked whether the fluorescent signal could be visualized through the eggshell in live embryos. Visualizing embryogenesis in living animals through the thick, semi-opaque eggshell and optically occlusive yolk has been a challenge in previous studies of *O. fasciatus* (Auman and Chipman, 2018; Panfilio, 2009; Panfilio et al., 2006). We focused specifically on katatrepsis, an essential embryonic movement (Panfilio, 2008). We found that all nuclei of the developing embryo were easily visualized through the eggshell, allowing us to capture the entire process of katatrepsis and appendage elongation over 36 hours (Fig. 4D and Supplementary Videos 1–3).

### Evidence for functionality of the Q system in *O. fasciatus*

To investigate whether this transgenesis method enables integration of large DNA payloads, we prepared the donor plasmid *pXL[Ofinv-QF2, 15×QUAS-mScarlet3, 9×P3-EGFP-SV40T]* (7,161 bp). This construct employs the Q system, a binary expression system composed of the transcriptional activator QF2 (a modified version of QF) and its binding site 15×QUAS (Potter et al., 2010; Riabinina et al., 2015) together with a 2.4 kb region upstream of the segment polarity gene *Of-invected* (*Of-inv*) (Accession No. OFAS025056), and the 9×P3 promoter-driven EGFP transformation marker (Fig. 6A). We performed three independent injections of the donor plasmid together with 1×NLS *hyPBase* mRNA. Among the surviving G_0_ nymphs, 50.0% (24/48), 26.1% (29/111), and 16.4% (18/110) showed mScarlet3 expression on their bodies at the nymphal instar stage (Table 2, Fig. 5B–F, and Supplementary Fig. 9). Notably, one of these 24 individuals displayed intense expression that partly appeared to align with thoracic segment boundaries. This individual also exhibited a segmental fusion phenotype (#1; Fig. 5B), which we speculate could be due to the strong activity of the *Of-inv* 2.4 kb regulatory region. At later nymphal stages, we observed strong mScarlet3 expression along the lateral edge of thoracic segments in a second individual (#2; Fig. 5C), broad mScarlet3 expression across the dorsal abdomen in a third (#3; Fig. 5D), and small patches and dots of expression in others (Supplementary Fig. 9). In individual #3 (Fig. 5D), clear segmental stripes of mScarlet3 expression were detected in the ventral abdomen (Fig. 5D), suggesting that *Of-inv* is expressed in post-embryonic development. Unfortunately, we did not achieve germ line transmission to the next generation, because animals with intense mScarlet3 expression died early and without producing progeny. To obtain information about whether the transgene had integrated into the genome despite this limitation, we performed splinkerette PCR on two G_0_ individuals (#4; Fig. 5E; #5; Fig. 5F-G). We detected multiple integration events in these individuals, indicating that our approach can mediate genomic integration of a large payload at least in G_0_ animals.

**Figure 5.**
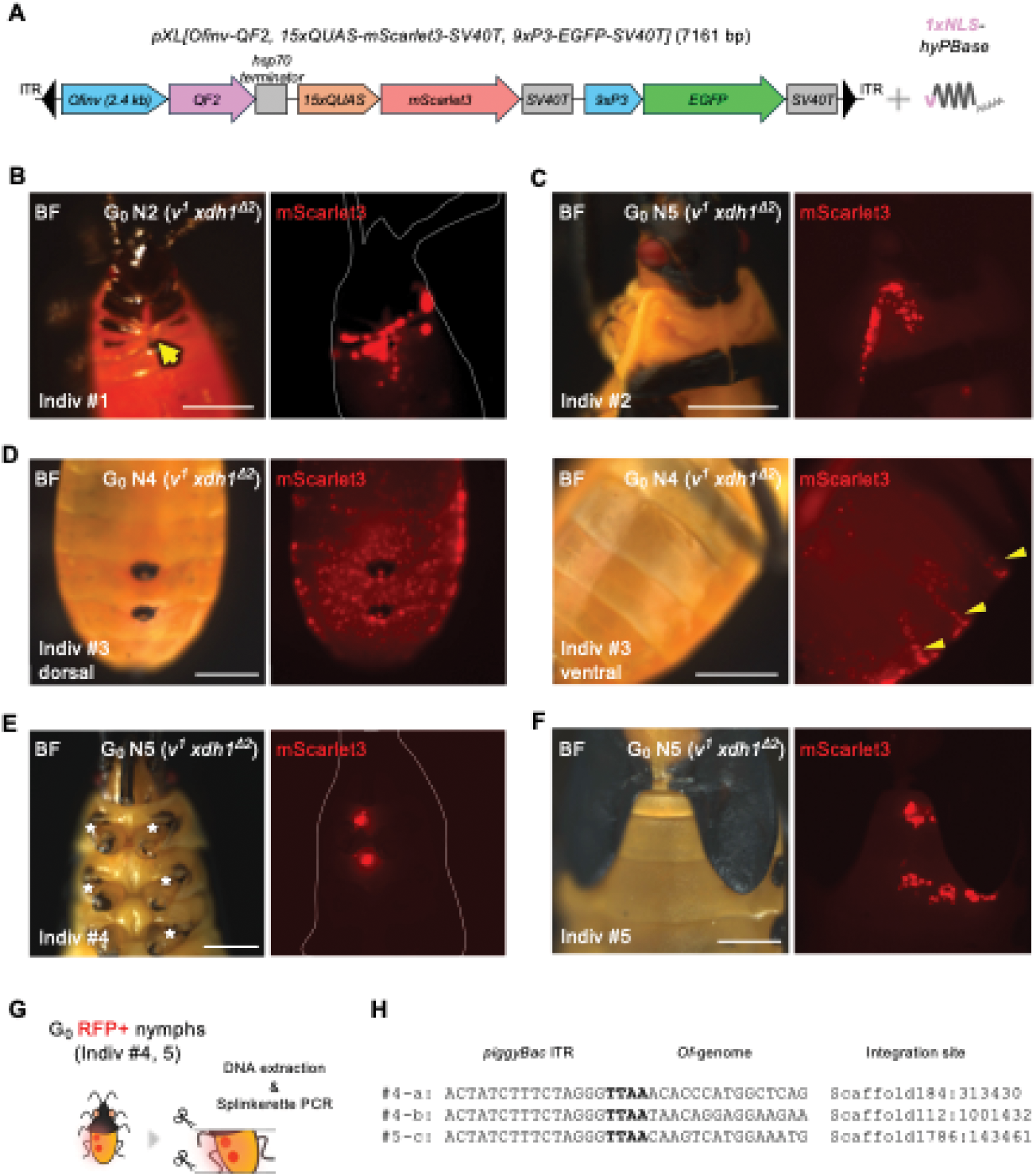
Proof of principle of the Q system using 1×NLS *hyPBase* in *O. fasciatus*. (A) Schematic of the *piggyBac* donor vector *pXL[Ofinv-QF2, 15×QUAS-mScarlet3-SV40T, 9×P3-EGFP-SV40T]* and *hyPBase* mRNA carrying 1×NLS. (B–F) Representative images of G_0_ mScarlet3-positive nymphs. Yellow arrow indicates the putative segmentation defect. Yellow arrowheads indicate the segmentally iterated abdominal expression of mScarlet3. Scale bars, 500 μm (B) and 1 mm (C–F). Anterior is up in (B–F). White asterisks in (E) indicate where legs were removed to improvevisibility of mScarlet3 expression. (G) Schematic of the splinkerette PCR procedure in G_0_ individuals. (H) Sequences of the junctions between the transgene and the flanking genomic DNA, determined by splinkerette PCR in G_0_ individuals.

## Discussion

We have established a system for efficient *piggyBac*-mediated transgenesis in the milkweed bug *O. fasciatus*. We show that efficiency is improved by the incorporation of NLS sequences into the hyPBase transposase. We also created a double mutant strain that facilitates more efficient screening with fluorescent 9×P3-driven transformation markers. This efficient and practical system enabled us to introduce multiple transgenic constructs carrying expression cassettes ranging from 1.8 to 7.2 kb in size into the *O. fasciatus* genome. Furthermore, we demonstrated proof of principle feasibility of the Q system-mediated binary expression system, expanding the genetic toolkit available in *O. fasciatus*.

CRISPR-mediated mutagenesis enabled us to revisit a classic study for the first time in half a century. Lawrence (1970) reported eye- and body-color mutants induced by ethyl methanesulfonate (EMS) treatment during the golden age of *O. fasciatus* research. Based on the descriptions of the phenotypes of these mutants (1970) and other studies (Liu, 2016; Reding et al., 2023), we focused on manipulating genes belonging to the pteridine pathway to generate a body-color mutant, and successfully established the *v^1^ xdh1*^Δ*2*^ double mutant strain showing body- and eye-color defects (Fig. 1). Although a recent study using RNAi-mediated functional analysis did not reveal a role for *xdh1* in eye-color formation (Reding et al., 2023), our results provide evidence that *xdh1* is involved in both body- and eye-color formation (Fig. 1). This discrepancy may be attributable to the often transient and incomplete nature of RNAi-mediated knockdown compared to a genomic loss-of-function allele. We further showed that both mScarlet-I and EGFP expressions can be detected through the eyes of the *v^1^ xdh1*^Δ*2*^ strain (Figs. 2, 3). Importantly, we have maintained this strain for over two years under the same conditions as the wild type strain and have observed no obvious developmental defects. As we have shown here, this strain is also used routinely for microinjection without noticeable problems. Together, these properties make the *v^1^ xdh1*^Δ*2*^ strain useful for developing future methods for transgenesis and targeted genomic integration, including CRISPR/Cas9-based knock-in (Matsuoka et al., 2025).

Another key finding is that we enhanced the overall efficiency of *piggyBac*-mediated transgenesis in *O. fasciatus* by incorporating SV40T NLS sequences into hyPBase. We found that 1× and 3×NLS (but not 5×NLS) increased both average STR and GTE compared to 0×NLS in *O. fasciatus* (Fig. 2), even though *piggyBac* transposase itself carries an intrinsic nuclear localization signal (Keith et al., 2008). We therefore used 1× and 3×NLS *hyPBase* mRNA for the subsequent transgenesis experiments. Copy-number estimates from G_1_ crossing experiments suggested that most transgenic lines generated with either 1× or 3×NLS *hyPBase* carried a single insertion (Supplementary Tables 5 and 7–8). The exception was the minimal donor plasmid *pXL[9×P3-EGFP-SV40T, attP]*, which led to multiple insertions (Supplementary Table 5). Since this plasmid is considerably smaller than the other transgenes and was tested only with 3×NLS *hyPBase*, we cannot distinguish the effect of NLS number from that of cargo size. Based on these results, either 1× or 3×NLS *hyPBase* can be used. We recommend 1×NLS *hyPBase* as a starting point, particularly when single-copy integration is desirable, with 3×NLS *hyPBase* as an alternative. A similar nuclear targeting approach may also be usefully applied to future phiC31-mediated site-specific integration in *O. fasciatus*, as a previous study demonstrated that the efficiency of phiC31 integrase is markedly improved by C-terminal incorporation of NLS in cultured mammalian cells (Andreas et al., 2002).

Interestingly, our results showed that 5×NLS completely abolished the transgenesis activity with no positive animals recovered, suggesting that an excessive number of NLS may be detrimental to transposase function in *O. fasciatus*. By contrast, all NLS-tagged *hyPBase* mRNAs functioned equally efficiently in *G. bimaculatus* (Supplementary Fig. 4), indicating that this nuclear targeting approach did not provide additional benefit in this species. We speculate that this species-specific difference may reflect differences in the timing of cellularization and/or biochemical properties between the two species. It would be interesting for future studies to investigate species-specific activity profiles of NLS-tagged *hyPBase* mRNAs across a broad range of insect species.

Identifying a suitable endogenous promoter driving ubiquitous and constitutive expression can be challenging in non-traditional insect models. Here we found that a promoter (*Gbact*) that drives ubiquitous, constitutive expression in a cricket species (Nakamura et al., 2010) did not drive such expression in *O. fasciatus*. Instead, the cricket promoter appeared to drive tissue-specific expression patterns, possibly due to capture by an endogenous *O. fasciatus* enhancer (Supplementary Fig. 7). In this case, the *Gbact* promoter appeared to function as a basal promoter. Similarly, in the case of GAP43-mScarlet3 driven by *Gbact*, we observed no clear membrane signals in ovaries or testes across multiple insertions (Supplementary Fig. 7). Although we cannot formally exclude the possibility that these expression patterns reflect position effects at the specific insertion sites, the consistent absence of ubiquitous expression across multiple independent insertions suggests that the *Gbact* promoter itself may not drive sufficiently strong ubiquitous expression in *O. fasciatus* to overcome the influence of nearby *O. fasciatus* enhancers. This observation is similar to one reported previously for the beetle *Tribolium castaneum*, wherein species-specific promoters were required to drive the desired expression patterns in transgenic animals (Schinko et al., 2010). To address this challenge, we identified the endogenous *O. fasciatus Act5c* promoter as one able to drive ubiquitous, constitutive expression and used it to successfully establish multiple histone-labelled transgenic strains (Fig. 4). Together, these transgenic strains and our exploration of heterologous and endogenous promoters provide valuable genetic and experimental resources for future work, facilitating comparative developmental studies in *O. fasciatus*.

Finally, we tested the feasibility of the Q system, a binary expression system, in *O. fasciatus* using a single-vector design, in which QF2 is driven by the 2.4-kb *Of-inv* upstream region and 15×QUAS-mScarlet3 is placed downstream of a transcriptional terminator (Fig. 5A). Across three independent injections, 16.4–50.0% of surviving G_0_ nymphs showed mScarlet3 expression, including segmentally patterned signal consistent with the expression of a segment polarity gene (Fig. 5B–F). Together with the detection of integration events in G_0_ individuals, these results indicate that a Q system-based transgene cassette exceeding 7 kb can be integrated and can drive reporter expression in *O. fasciatus*. This may provide a platform for amplifying and visualizing cis-regulatory element activity in future studies. However, although the terminator is expected to minimize transcriptional readthrough from the QF2 cassette, our single-vector design cannot formally exclude the possibility that mScarlet3 expression arises from transcriptional readthrough, or from direct activation of the reporter’s minimal promoter by the *Of-inv* regulatory region. Future work, including crossing QF2 driver and QUAS reporter lines, will be needed to provide definitive validation. Notably, one of the G_0_ nymphs we obtained with strong transgene expression exhibited a segmental fusion phenotype (Fig. 5), which we speculate could be due to excessive *Of-inv* promoter activity; this animal died before late nymphal stages. This could mean that maintaining moderate transcriptional expression levels of some constructs may be important for viability.

Taken together, our results have substantially expanded the functional genetic toolkit of *O. fasciatus* as a comparative model insect for Evo-Devo research. By combining a visibly light-colored mutant background, NLS-tagged hyPBase activity, and the Q system-based binary expression platform, we have established a versatile and practical framework for functional genetics in this species. These technological advances position *O. fasciatus* as an increasingly powerful system for addressing fundamental biological questions in development and evolution. Moreover, the nuclear targeting approach for improving overall transgenesis efficiency will offer a scalable framework that may benefit other non-model organisms, broadening the scope of comparative genetics in Evo-Devo studies beyond established model species.

## Materials and Methods

### Animal husbandry

*Oncopeltus fasciatus* was maintained on sunflower seeds at 28–30 °C and 60% relative humidity with 12L:12D as described previously (Ewen-Campen et al., 2011). *v^1^* mutant *O. fasciatus* were kindly provided by Katie Reding and Leslie Pick (University of Maryland, College Park, MD, USA). A *Gryllus bimaculatus* colony derived from the *gwhite* strain at Tokushima University (Niwa et al., 1997) was maintained on cat food (Purina) under identical environmental conditions.

### Embryo microinjection

Injection needles were prepared by pulling glass capillaries (Cat#1B100-4, World Precision Instruments) with a Sutter P-97 Flaming/Brown micropipette puller. In *O. fasciatus*, eggs were collected within four hours after egg laying and injections were completed within two hours after egg collection. Two glass microscope slides were taped together with double-sided tape to create a ledge with exposed double-sided tape, against which embryos were aligned. Eggs were submerged in tap water for 20-40 minutes to reduce the inner pressure for ease of injection. Injected eggs were kept at 28 ℃. In *G. bimaculatus*, eggs were collected within two hours after egg laying and injection procedures were completed within two hours after egg collection. Eggs were aligned in injection plates with wells made of 2% agarose dissolved in MilliQ water. Injected eggs were kept at 28℃ until the desired stage.

### CRISPR-Cas9-mediated targeted mutagenesis

Alt-R S.p. Cas9 Nuclease V3 (Cat# 1081059) and tracrRNA (Cat# 1072533) were chemically synthesized by Integrated DNA Technologies (IDT). *Of-xdh1* crRNA was designed to target the coding sequence using CRISPR Optimal Target Finder (Gratz et al., 2014). Synthetic crRNA was provided by IDT. Cas9 ribonucleoprotein complex (RNP) was prepared according to IDT instructions. Briefly, tracrRNA and crRNA were resuspended to 100 μM in Nuclease-Free Duplex Buffer (Cat# 11-01-03-01). 5 μl of each RNA were mixed, heated to 95℃ for five minutes, and allowed to cool to room temperature to form 50 μM gRNA duplex. 3 μl gRNA duplex and 2 μl Cas9 protein were mixed and incubated at room temperature for ten minutes to form Cas9/gRNA RNP at roughly 25 μM. Finally, RNP was diluted tenfold in 1×PBS with 0.05% phenol red.

### Donor plasmid construction

The *pXL[Gbact-H2B-EGFP]* vector was kindly provided by Taro Nakamura (National Institute of Basic Biology, Okazaki, Japan) and used as a backbone vector for all *piggyBac*-mediated transgenesis in this study. The H2B CDS, derived from *G. bimaculatus*, was used as a nuclear marker. The plasmid encoding an insect codon-optimized variant of mScarlet-I, *pUC[9×P3-mScarlet-I-SV40T]*, was kindly provided by Takaaki Daimon (Kyoto University, Kyoto, Japan). Using these constructs, the donor plasmids *pXL[9×P3-mScarlet-I-SV40T]* and *pXL[9×P3-EGFP-SV40T, attP]* were assembled. To generate *pXL[Gbact-H2B-mScarlet3-SV40T]* and *pXL[Gbact-GAP43-mScarlet3-SV40T, 9×P3-EGFP-SV40T]*, the plasmid *5_T7-GAP43-mScarlet3* (Addgene #225932) was used in addition to the plasmids described above. To generate *pXL[Ofact-H2B-mScarlet3-SV40T]*, the 2.5 kb region upstream of *Of-act5ca* was amplified from *O. fasciatus* genomic DNA, and other sequences were obtained from *pXL[Gbact-H2B-mScarlet3-SV40T].* To generate *pXL[Ofinv-QF2, 15×QUAS, 9×P3-EGFP]*, the 2.4 kb region upstream of *Of-inv* was amplified from *O. fasciatus* genomic DNA, and QF2 and QUAS sequences were obtained from plasmids *pBAC-DsRed-QF2-hsp70* (Addgene #104876) and *PBAC-ECFP-15×QUAS_TATA-SV40* (Addgene #104875), respectively. Nucleotide sequences of all donor plasmids are provided in Supplementary Data 1. For microinjection, donor plasmids were diluted to 200 ng/μL in nuclease-free water.

### mRNA preparation

The expression vector for mammalian codon-optimized hyPBase (*pCMV-hyPBase*) (Yusa et al., 2011) was kindly provided by Takahiro Ohde (Kyoto University, Kyoto, Japan). To generate *pCMV-3×NLS-hyPBase*, the backbone of *pCMV-hyPBase* was linearized by inverse PCR using primers partially encoding the SV40 NLS. The remaining NLS sequences were provided as annealed oligonucleotides, and the fragments were assembled using the Gibson Assembly Cloning Kit (New England Biolabs, Cat#E5510S), so that three tandem SV40 NLS repeats were fused to the N-terminus of hyPBase. *pCMV-1×NLS-hyPBase* and *pCMV-5×NLS-hyPBase* were generated from *pCMV-3×NLS-hyPBase* using the Q5 Site-Directed Mutagenesis Kit (New England Biolabs, Cat#E0554S) according to the manufacturer’s protocol. Amino acid sequences of all hyPBase variants are provided in Supplementary Data 2, and the primers and oligonucleotides used are listed in Supplementary Table 9. For *hyPBase* mRNA synthesis, the plasmids were linearized with BbsI (New England Biolabs, Cat#R0539S), and capped mRNA was transcribed using the mMESSAGE mMACHINE T7 Transcription kit (Thermo Fisher Scientific, Cat#AM1344), followed by lithium chloride purification. The concentration of purified mRNA was determined by measuring absorbance at 260 nm using a NanoDrop Eight spectrophotometer (Thermo Fisher Scientific). For microinjection, purified mRNAs were diluted to 400 ng/μL in nuclease-free water.

### Genomic DNA extraction and genotyping

For genotyping, genomic DNA was extracted from a single leg of animals. Legs were homogenized in squish buffer (10 mM Tris-HCl pH 7.5, 1 mM EDTA, 25 mM NaCl, 200 μg/ml Proteinase K), incubated at 60℃ for two hours, and then at 95℃ for three minutes to inactivate Proteinase K. Genomic PCR was performed using Phusion HF DNA polymerase (New England Biolabs Cat#M0530L). Targeted mutagenesis was detected using the Alt-R Genome Editing Detection Kit (IDT, Cat#1075932), and fixed alleles were confirmed by Sanger sequencing. For splinkerette PCR, genomic DNA was prepared using the DNeasy & Tissue kit (QIAGEN, Cat#69506).

### Splinkerette PCR

Genomic DNA was extracted from two G_2_ animals carrying *9×P3-EGFP-SV40T, attP*, two G_2_ animals carrying *Gbact-H2B-mScarlet3-SV40T*, and two G_0_ animals showing mScarlet3 expression from *Ofinv-QF2, 15×QUAS-mScarlet3, 9×P3-EGFP-SV40T*. Extracted genomic DNA was digested with BstYI (New England Biolabs, Cat#R0523S), followed by self-ligations. Two rounds of splinkerette PCR were performed using KOD-One (TOYOBO, Cat#KMM-101NV). All procedures for splinkerette PCR were performed as described previously (Potter and Luo, 2010). For G_2_ animals, the PCR products were purified and directly sequenced by Sanger sequencing. For G_0_ animals, since the PCR products yielded faint, smeared bands, they were cloned into the pGEM-T easy Vector (Promega, Cat#A1360). Primers are listed in Supplementary Table S9.

### Screening transgenic animals

Animals were anesthetized with CO_2_ and screened for fluorescence using a Zeiss Axio Zoom V16 stereomicroscope. Images were captured using a Zeiss Axical 512 color camera or HAMAMATSU C11440-22CU digital camera.

### Nuclear and F-actin staining

Ovaries were dissected from transgenic G_1_ adults and fixed with 4% formaldehyde in 1× PBS for 30 minutes at room temperature, followed by three washes with PBT (1× PBS with 0.1% Triton X-100). Samples were incubated with 5 ng/μl Hoechst 33342 and Alexa Fluor 488-phalloidin (1:200; Thermo Fisher Scientific, Cat#A12379) in PBT at room temperature for one hour or at 4℃ overnight. Samples were washed four times over one hour with PBT. After washing, samples were mounted in 100% glycerol. Images were taken using a Nikon CSU-W1 spinning disk inverted confocal microscope.

### Live imaging

Appropriately staged eggs were collected under a Zeiss Axio Zoom V16 stereomicroscope and attached to a standard 90 mm polystyrene Petri dish using heptane glue. Live imaging was performed on a Nikon CSU-W1 spinning disk inverted confocal microscope with 10× air objective at 29□. Image stacks comprising approximately 40 optical sections (4 μm z-steps) were taken at 10 minute intervals.

### Use of artificial intelligence tools

The authors used ChatGPT (OpenAI) and Claude (Anthropic) to improve the clarity and readability of the text. All authors reviewed and edited all output and take responsibility for all content. No AI tools were used to generate original data, results, or scientific content.

## Acknowledgements

We thank Tarun Kumar and Daniella Haber for helping with animal husbandry, Suhrid Ghosh for assistance with live imaging, and all members of the Extavour lab for discussion. Y.S. is supported by the Human Frontier Science Program (HFSP) Fellowship LT0048/2025-L (DOI: https://doi.org/10.52044/HFSP.LT00482025-L.pc.gr.231571). Y.S. was a Japan Society for the Promotion of Science (JSPS) Overseas Research Fellow. This study was supported by NSF award IOS-2220747, NIH award 1R01GM143611, and funds from the Howard Hughes Medical Institute (HHMI) to C.G.E., who is an HHMI investigator.

## Author contributions

Conceptualization: Y.S.; Methodology: Y.S.; Validation: Y.S.; Formal analysis: Y.S.; Investigation: Y.S., Y.H., A.W., J.A.K.; Resources: Y.S., Y.H., A.W., J.A.K.; Data curation: Y.S.; Writing - original draft: Y.S.; Writing - review & editing: Y.S., Y.H., A.W., J.A.K., C.G.E.; Visualization: Y.S.: Supervision: Y.S., C.G.E.; Project administration: Y.S., C.G.E.; Funding acquisition: Y.S., C.G.E.

## Data and resource availability

All data and resources in this paper will be shared upon request.

