## Supplemental Information for "A nuclear targeting approach enables efficient transgenesis in the milkweed bug *Oncopeltus fasciatus*"

This file contains:

SI Figures S1-S9

SI Tables S1-S9

SI Data 1 and Data 2

SI Video Legends

SI References

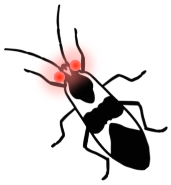

### Supplementary Figures

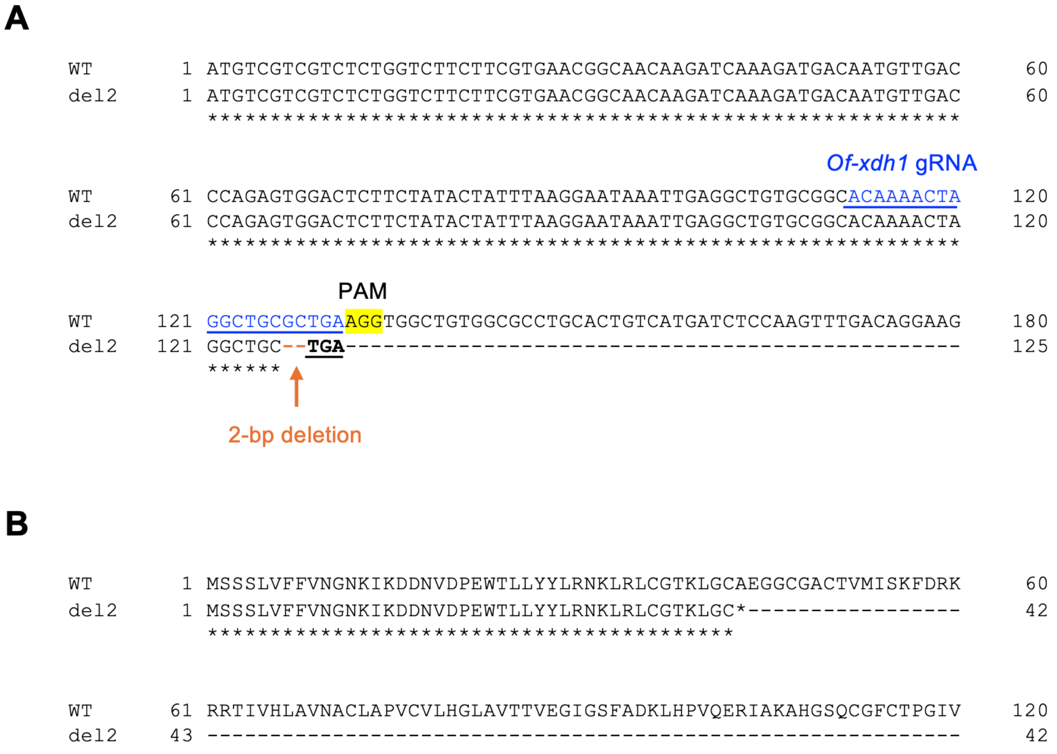


**Figure S1. Nucleotide and amino acid sequence of *Of-xdh1* in the *xdh1*$^{\Delta 2}$ mutant.** (A) Alignment of the nucleotide sequences of *Of-xdh1* around the gRNA target site of WT and *xdh*$^{\Delta 2}$ strains. Note the presence of a 2-bp deletion in the *xdh1*$^{\Delta 2}$ allele (shown in orange). The premature stop codon induced by this deletion is indicated in bold black. (B) Alignment of the amino acid sequences of *Of-xdh1* of the WT and *xdh1*$^{\Delta 2}$ strains. The 2-bp deletion caused a frameshift mutation in the *xdh1*$^{\Delta 2}$ allele.

**A**

```
WT      1 ATGGAGCCCGACAGCAATTGCAGATCTTCCGAGATCCAGAA-ATGTGGAGAGCAAAGTGG     59
v1      1 ATGGAGCCCGACAGCAATTGCAGATCTTCCGAGATCCAGAAGATGTGGAGAGCAAAGTGG     60
          ***************************************** ******************

                                          1-bp insertion

WT     60 AATGCTGTATGGGGATTACCTCCAACTTGACAAAATACTGAAGGCCCAAAGATTGCTGAG    119
v1     61 AATGCTGTATGGGGATTACCTCCAACTTGA------------------------------     90
          ******************************
```

**B**

```
WT      1 MEPDSNCRSSEIQKCGEQSGMLYGDYLQLDKILKAQRLLSESNNATVMDEHLFIVTHQGN     60
v1      1 MEPDSNCRSSEIQKMWRAKWNAVWGLPPT*------------------------------     29
          **************

WT     61 KLNSTQEMTNEGLNERQTLDVVKRLNRIVLILKLLVEQVMILETMTPLDFMEFRDYLCPA    120
v1     29 ------------------------------------------------------------     29

WT    121 SGFQSAQFRLIENKLGVKTEHRVRSFQHYSTVFERDKEALNEIKKSETEQSLADLIEQWL    180
v1     29 ------------------------------------------------------------     29

WT    181 ERTPGLDENGFDFWGKYIKNVEFLLQHYYEKAMKSNSETEKTTKLAEYEKKKTIFNSVLN    240
v1     29 ------------------------------------------------------------     29

WT    241 KDMHDLFVQRGERRFTHKAFQGAIMITFYRDEPRFSQPHQILTLLMDIDSLVTKWRLCSK    300
v1     29 ------------------------------------------------------------     29

WT    301 TWINYYNDKYKVLLISDRYKCFLDLFNLSSFLLPRNFIPPLSSSVRNQINIHTWNTKKQT    360
v1     29 ------------------------------------------------------------     29

WT    361 EDHEED*                                                          366
v1     29 ------                                                            29
```

**Figure S2. Nucleotide and amino acid sequence of *Of-v* in the *v*$^1$ mutant.** (A) Alignment of the nucleotide sequences of *Of-v* around the gRNA target site of wild type (WT) and *v*$^1$ strains. Note the presence of a 1-bp insertion in the *v*$^1$ allele (shown in red). The premature stop codon induced by this insertion is indicated in bold black. Further details are described in (Reding and Pick, 2023). (B) Alignment of the amino acid sequences of *Of-v* of the WT and *v*$^1$ strains. The 1-bp insertion caused a frameshift mutation in the *v*$^1$ allele (shown in red).

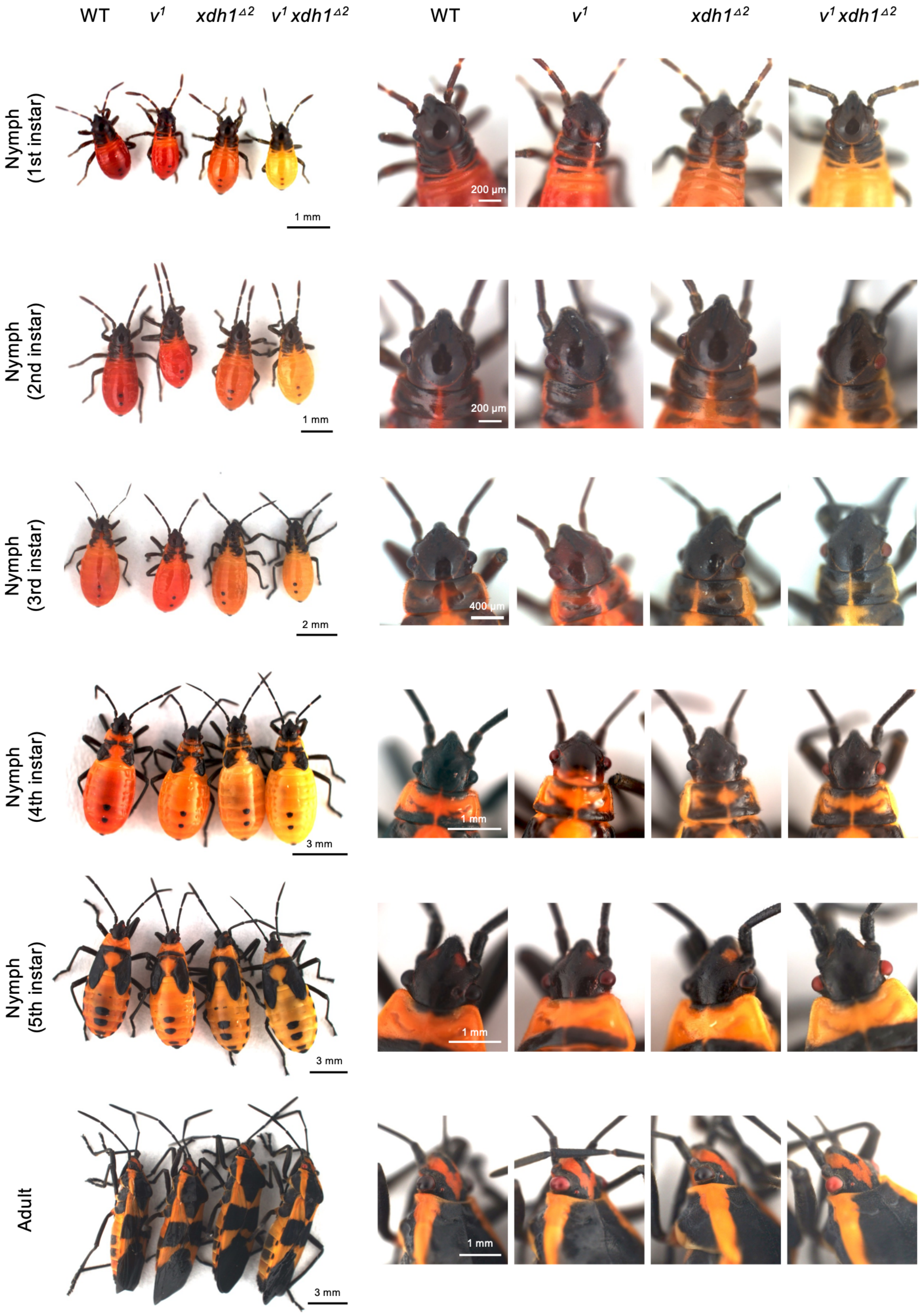


**Figure S3. Comparisons of phenotypes of wild type (WT), $v^1$, $xdh1^{\Delta2}$, and $v^1$ $xdh1^{\Delta2}$ throughout postembryonic development.** All images of nymphs and close-up views (bottom right) are shown in dorsal view; low-magnification adult image (bottom left) is shown in lateral view. Anterior is up in all images.

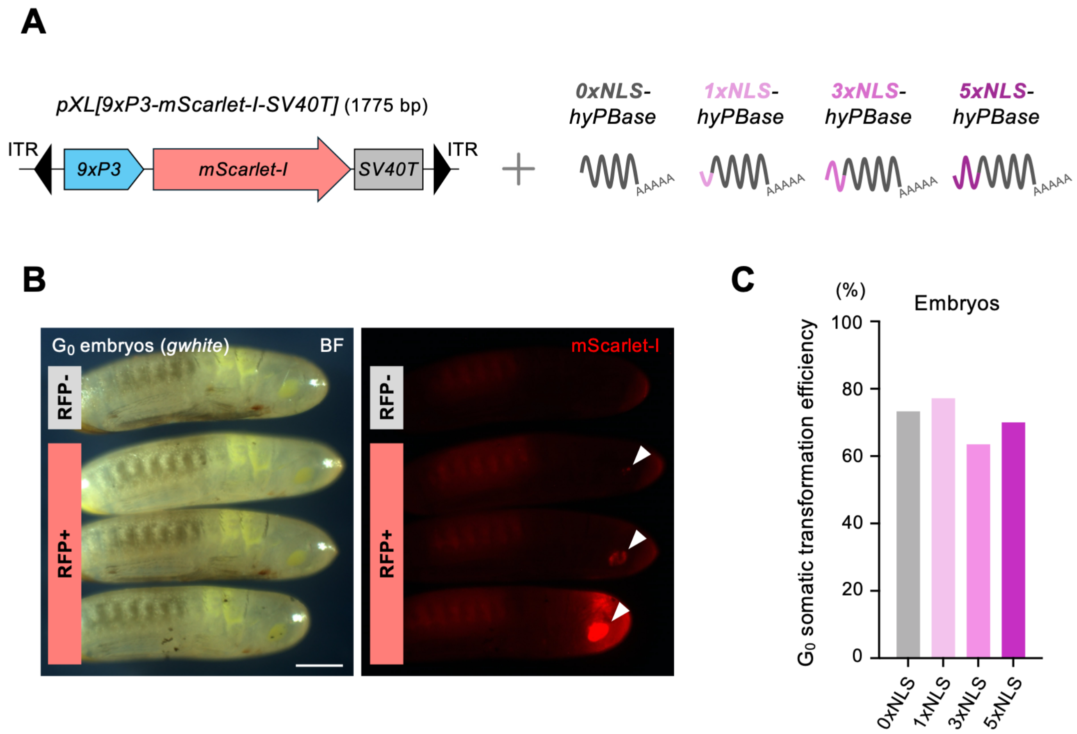


**Figure S4. Investigation of a nuclear targeting approach for *piggyBac*-mediated transgenesis in *G. bimaculatus*.** (A) Schematic of the *piggyBac* donor vector *pXL[9×P3-mScarlet-I-SV40T]* and *hyPBase* mRNA carrying 0×, 1×, 3×, or 5× NLS. (B) Representative images of $G_0$ RFP-positive embryos (embryonic stage >13) (Donoughe and Extavour, 2016). Scale bar, 500 µm. Arrowheads indicate eyes. Anterior is to the right. (C) $G_0$ somatic transformation rate (STR) across 0×, 1×, 3×, or 5×NLS *hyPBase* conditions.

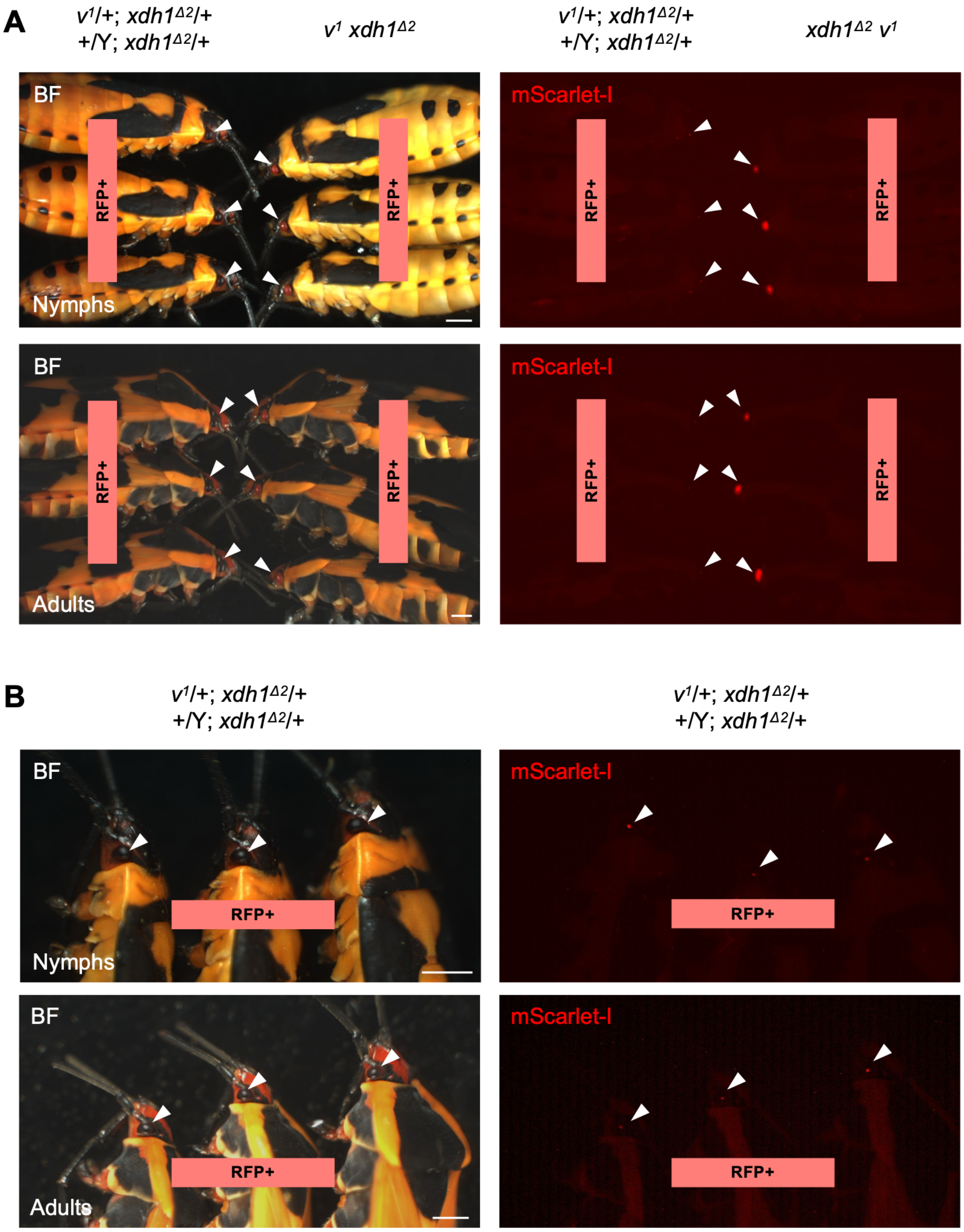


**Figure S5. Comparison of mScarlet-I detection between wild type and $v^1$ $xdh1^{\Delta 2}$ backgrounds in *O. fasciatus*.** (A) Representative images of mScarlet-I-positive nymphs and adults in phenotypically wild-type backgrounds ($v^1$/+; $xdh1^{\Delta 2}$/+ or +/Y; $xdh1^{\Delta 2}$/+) and the $xdh1^{\Delta 2}$ $v^1$ mutant background. Arrowheads indicate eyes. (B) Enlarged images of mScarlet-I-positive nymphs and adults in the wild type background ($xdh1^{\Delta 2}$/+ $v^1$/+ or $xdh1^{\Delta 2}$/+ +/Y). Scale bar, 1 mm. Transgene copy number was not controlled, as this comparison was intended to assess the visibility of the eye marker.

#### Integration site in 9xP3-EGFP, attP strain 5A

TATTTCAAGAATGCATGCGTCAATTTTACGCAGACTATCTTTCTAGGG**TTAA**TTAAAATTGCGTGGTACGATTTG
TAATCTACAGTTTACCCGGACACTAAATTACAATTTTACGTTTTATTGGCCGAAAATAGTTGTTTTTCTAGACAA
AA

Scaffold407: 476763-476862 (pident = 100%)

*The transgene was inserted into an intron of OfasTmpM006116-RA

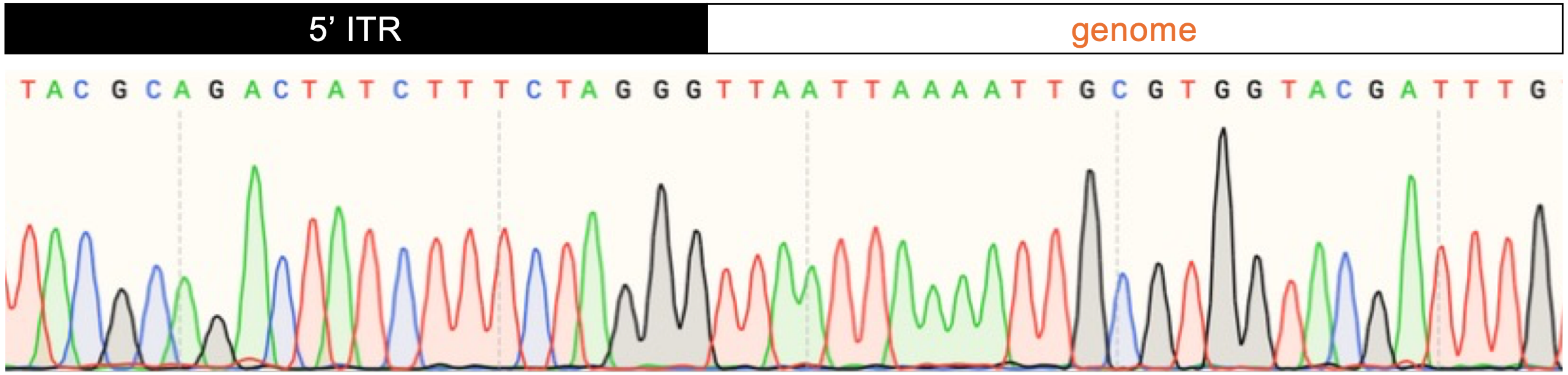


**Figure S6. Nucleotide sequences around the transgene integration site in the *9×P3-EGFP*, *attP* strain in *O. fasciatus* as determine by splinkerette PCR.**

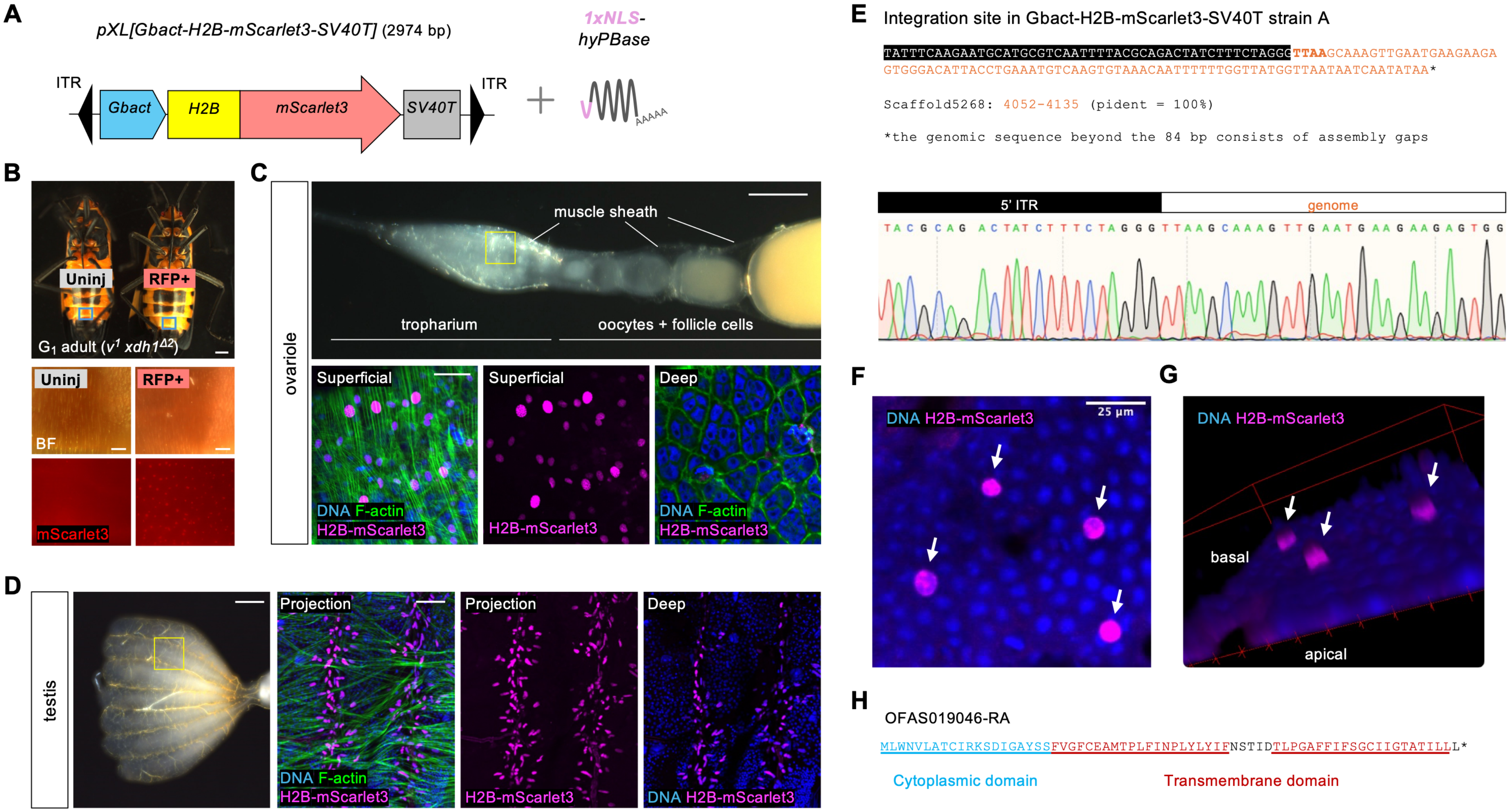
A
pXL[Gbact-H2B-mScarlet3-SV40T] (2974 bp)
1xNLS-hyPBase
ITR
Gbact
H2B
mScarlet3
SV40T
ITR
AAAAA
B
Uninj
RFP+
G1 adult (v1 xdh1Δ2)
Uninj
RFP+
BF
mScarlet3
C
ovariole
muscle sheath
tropharium
oocytes + follicle cells
Superficial
Superficial
Deep
DNA F-actin
H2B-mScarlet3
H2B-mScarlet3
DNA F-actin
H2B-mScarlet3
D
testis
Projection
Projection
Deep
DNA F-actin
H2B-mScarlet3
H2B-mScarlet3
DNA H2B-mScarlet3
E
Integration site in Gbact-H2B-mScarlet3-SV40T strain A
TATTTCAAGAATGCATGCGTCAATTTTACGCAGACTATCTTTCTAGGGTTAAGCAAAGTTGAATGAAGAAGA
GTGGGACATTACCTGAAATGTCAAGTGTAAACAATTTTTTGGTTATGGTTAATAATCAATATAA*
Scaffold5268: 4052-4135 (pident = 100%)
*the genomic sequence beyond the 84 bp consists of assembly gaps
5' ITR
genome
F
DNA H2B-mScarlet3
25 µm
G
DNA H2B-mScarlet3
basal
apical
H
OFAS019046-RA
MLWNVLATCIRKSDIGAYSSFVGFCEAMTPLFINPLYLYIFNSTIDTLPGAFFIFSGCIIGTATILLL*
Cytoplasmic domain
Transmembrane domain

**Fig S7 (previous page). Establishment of the tissue-specific histone-labelled strain using 1×NLS *hyPBase* in *O. fasciatus*.** (A) Schematic of the *piggyBac* donor vector *pXL[Gbact-H2B-mScarlet3-SV40T]* and *hyPBase* mRNA carrying 1×NLS. (B) Representative images of $G_1$ mScarlet3-positive adults (ventral view). Anterior is up. Scale bars, 1 mm (whole-body image) and 100 µm (close-up images). (C) Representative brightfield image (top) and confocal images (bottom) of an ovariole from a $G_1$ transgenic adult, co-stained for DNA (blue) and F-actin (green). The yellow box in the brightfield image indicates the approximate region shown in the confocal images. Confocal images are single optical sections. In the superficial focal plane, H2B-mScarlet3 expression (magenta) is detectable in muscle sheath nuclei. In the deep focal plane, H2B-mScarlet3 expression is not detectable in the tropharium nuclei. Scale bars, 300 µm (brightfield image) and 50 µm (confocal images) (D) Representative brightfield image (left) and confocal images (right three panels) of a testis of $G_1$ transgenic adult testis co-stained for DNA (blue) and F-actin (green). H2B-mScarlet3 expression (magenta) was detected in muscle sheath nuclei, but not in sperm cells. Scale bars, 300 µm (brightfield image) and 50 µm (confocal images). Anterior is to the left in (C) and (D). (E) Nucleotide sequences around the transgene integration site in the *H2B-mScarlet3-SV40T* strain as determined by splinkerette PCR. (F) Confocal image of the abdominal integument of the *H2B-mScarlet3-SV40T* strain co-stained for DNA. Epidermal nuclei were labelled with blue, but not labelled with magenta (H2B-mScarlet3). Arrows indicate cells labelled with H2B-mScarlet3. (G) 3D reconstruction of the abdominal integument of the H2B-mScarlet3 strain co-stained for DNA, showing that the histone-labelled nuclei with H2B-mScarlet3 were located underneath the epidermal nuclei. Arrows indicate cells labelled with H2B-mScarlet3. (H) Amino acid sequences of the uncharacterized protein (OFAS019046-RA) encoded by the locus containing the transgene insertion site.

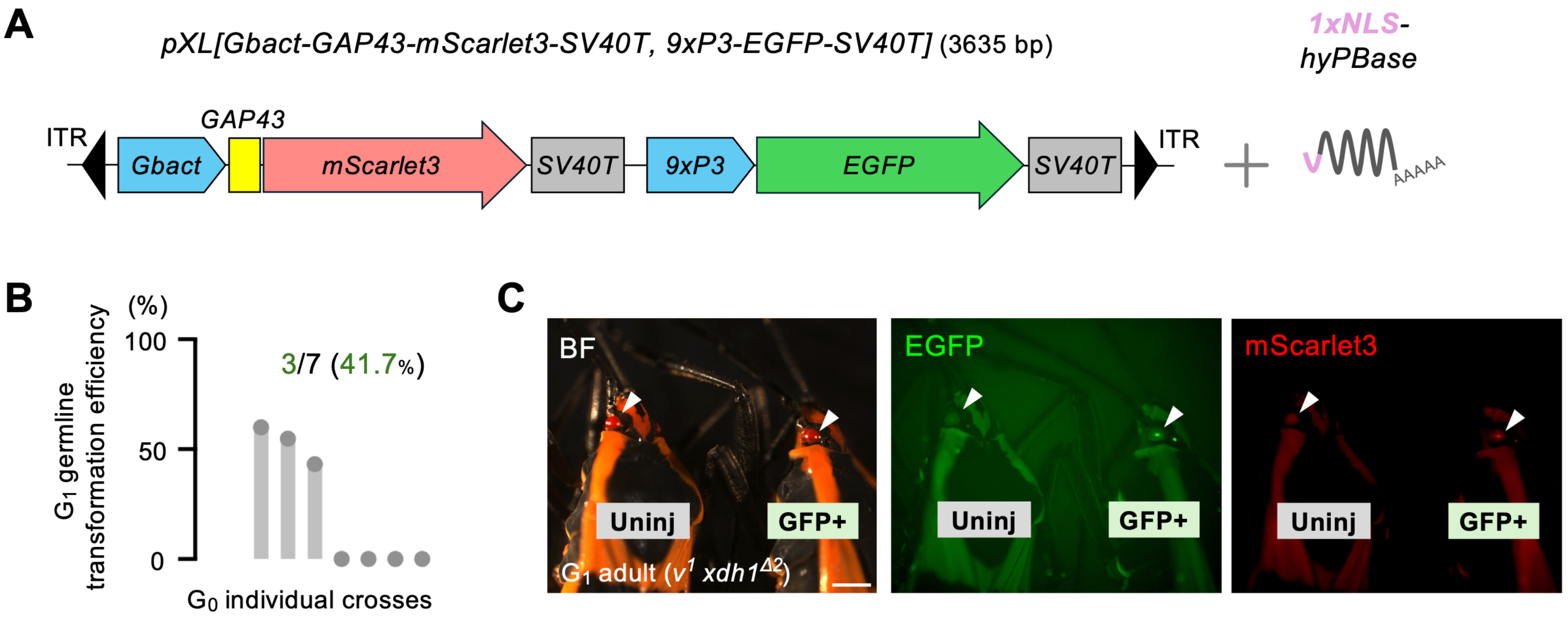


**Figure S8. Attempt to generate a membrane-labelled strain using 1×NLS *hyPBase* in *O. fasciatus*.** (A) Schematic of the *piggyBac* donor vector *pXL[Gbact-GAP43-mScarlet3-SV40T, 9xP3-EGFP-SV40T]* and *hyPBase* mRNA carrying 1×NLS. (B) $G_1$ germ line transformation efficiency (GTE). Each bar represents the proportion of EGFP-positive animals among $G_1$ progeny from an individual cross. Each dot represents an individual backcross that yielded $G_1$ progeny. Green numbers indicate the number of crosses yielding heritable transgenes and the corresponding percentages. (C) Representative images of $G_1$ EGFP-positive adults. Weak mScarlet3 expression in their eyes was also observed.

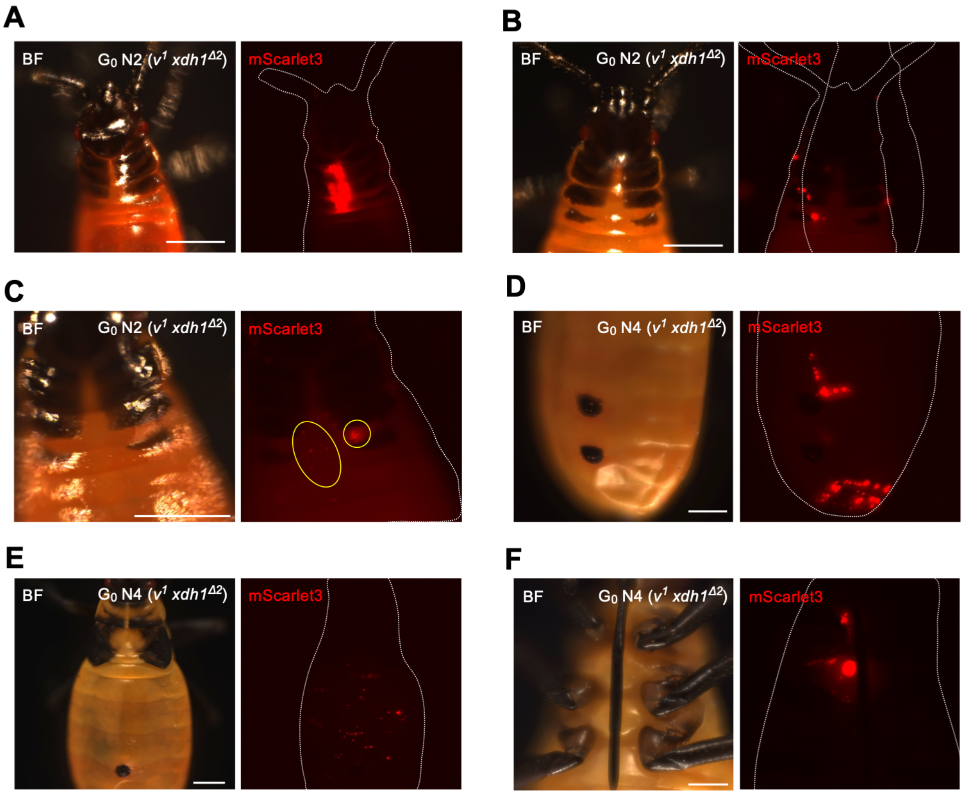


**G**

Integration sites in Ofinv-QF2, 15×QUAS-mScarlet3-SV40T, 9×P3-EGFP-SV40T, $G_0$ indiv#4

```
TATTTCAAGAATGCATGCGTCAATTTTACGCAGACTATCTTTCTAGGGTTAAACACCCATGGCTCAGGACAAGCTTC
TATATTATAAATAAAAGATTGACCTGGGCGGGATTAGAACCCGCGGCGTCAGGATTGCGAGACCTGTACGGATCC

Scaffold184: 313430-313529 (pident = 100%)


TATTTCAAGAATGCATGCGTCAATTTTACGCAGACTATCTTTCTAGGGTTAATAACAGGAGGAAGAATGAGAGAGTA
GTCAACACTCACAAACACACAAAATAGAAATAACGCAAAGTATTGCGAATATTTAGTTTTCTTTTTATATATATT

Scaffold112: 1001432-1001531 (pident = 100%)
```

**H**

Integration sites in Ofinv-QF2, 15×QUAS-mScarlet3-SV40T, 9×P3-EGFP-SV40T, $G_0$ indiv#5

```
TATTTCAAGAATGCATGCGTCAATTTTACGCAGACTATCTTTCTAGGGTTAACAAGTCATGGAAATGTGAACTTAGG
GACAGGACGTCCTCTCTCATCTTCATGATAGCCCTGAAGATGGCTTGCAATGTAAGCCGAAACGTTGGCAGATAA

Scaffold1786: 143362-143461 (pident = 99%)
```

**Figure S9. Additional data on Q system application using 1×NLS *hyPBase* in *O. fasciatus*.** (A–F) Representative images of $G_0$ mScarlet3-positive nymphs. White dotted lines indicate animal boundaries. Yellow dotted circles indicate mScarlet3 expression. Scale bars, 500 µm. Anterior is up in (A–F). (G-H) Nucleotide sequences around the transgene integration site of animals in the injected generation as determined by splinkerette PCR.

#### Supplementary Tables

**Table S1. Detailed results of piggyBac-mediated transgenesis experiments with hyPBase mRNA carrying 0×, 1×, 3×, or 5×NLS in *G. bimaculatus*.** Each row shows the results of an independent experiment. Percentages shown in parentheses indicate somatic transformation rates (STR).

| Donor plasmid | Transposase (mRNA) | Injected eggs | % of injected eggs survived to screening | Screened G0 embryos | mScarlet-I **+** nymphs | mScarlet-I **-** nymphs |
|---|---|---|---|---|---|---|
| pXL[9xP3-mScarlet-I-SV40T] | 0xNLS-hyPBase | 194 | 69.6% | 135 | 99 (73.3%) | 36 |
| pXL[9xP3-mScarlet-I-SV40T] | 1xNLS-hyPBase | 194 | 70.1% | 136 | 105 (77.2%) | 31 |
| pXL[9xP3-mScarlet-I-SV40T] | 3xNLS-hyPBase | 190 | 72.1% | 137 | 87 (63.5%) | 50 |
| pXL[9xP3-mScarlet-I-SV40T] | 5xNLS-hyPBase | 192 | 67.7% | 130 | 91 (70.0%) | 39 |

**Table S2. Results of crossing experiments with 0×NLS-hyPBase.** F, female. M, male. GTR, germ line transformation rate.

| mRNA | donor plasmid |
|---|---|
| 0xNLS-hyPBase | 9xP3-mScarlet-I-SV40T |

| G0 indiv# | mScarlet-I expression | sex | crossed with | Screened G1 animals | RFP+ | RFP- | GTR (%) |
|---|---|---|---|---|---|---|---|
| 1 | + | M | $v^{1}$ $xdh1^{\Delta}$ | 60 | 0 | 60 | 0 |
| 2 | + | F | $v^{1}$ $xdh1^{\Delta}$ | 57 | 0 | 57 | 0 |
| 3 | + | M | $v^{1}$ $xdh1^{\Delta}$ | 57 | 0 | 57 | 0 |
| 4 | + | F | $v^{1}$ $xdh1^{\Delta}$ | 54 | 31 | 23 | 57.4 |
| 5 | + | F | $v^{1}$ $xdh1^{\Delta}$ | 57 | 0 | 57 | 0 |
| 6 | + | F | $v^{1}$ $xdh1^{\Delta}$ | 13 | 0 | 13 | 0 |

**Table S3. Results of crossing experiments with 1×NLS-hyPBase.** F, female. M, male. GTR, germ line transformation rate. Data from two independent injection experiments are shown.

| mRNA | donor plasmid | | | | | | |
|---|---|---|---|---|---|---|---|
| 1xNLS-hyPBase | 9xP3-mScarlet-I-SV40T | | | | | | |
| G0 indiv# | mScarlet-I expression | sex | crossed with | Screened G1 animals | RFP+ | RFP- | GTR (%) |
| 1 | + | F | $v^1$ $xdh1^{\Delta 2}$ | 55 | 0 | 55 | 0 |
| 2 | + | F | $v^1$ $xdh1^{\Delta 2}$ | 45 | 0 | 45 | 0 |
| 3 | + | F | $v^1$ $xdh1^{\Delta 2}$ | 64 | 0 | 64 | 0 |
| 4 | + | F | $v^1$ $xdh1^{\Delta 2}$ | 52 | 0 | 52 | 0 |
| 5 | + | F | $v^1$ $xdh1^{\Delta 2}$ | 53 | 44 | 9 | 83.0 |
| 6 | + | M | $v^1$ $xdh1^{\Delta 2}$ | sterile | 0 | 0 | NA |
| 7 | + | M | $v^1$ $xdh1^{\Delta 2}$ | 24 | 11 | 13 | 45.8 |
| 8 | + | M | $v^1$ $xdh1^{\Delta 2}$ | 50 | 0 | 50 | 0 |
| 9 | + | F | $v^1$ $xdh1^{\Delta 2}$ | 34 | 0 | 34 | 0 |
| 10 | + | F | $v^1$ $xdh1^{\Delta 2}$ | 43 | 0 | 43 | 0 |
| 11 | + | F | $v^1$ $xdh1^{\Delta 2}$ | 61 | 54 | 7 | 88.5 |
| 12 | + | M | $v^1$ $xdh1^{\Delta 2}$ | 39 | 18 | 21 | 46.2 |
| 13 | + | M | $v^1$ $xdh1^{\Delta 2}$ | 51 | 0 | 51 | 0 |
| 14 | + | M | $v^1$ $xdh1^{\Delta 2}$ | 52 | 35 | 17 | 67.3 |
| 15 | + | M | $v^1$ $xdh1^{\Delta 2}$ | 48 | 0 | 48 | 0 |
| 16 | + | M | $v^1$ $xdh1^{\Delta 2}$ | 26 | 0 | 26 | 0 |
| 17 | + | M | $v^1$ $xdh1^{\Delta 2}$ | 62 | 0 | 62 | 0 |
| 18 | + | M | $v^1$ $xdh1^{\Delta 2}$ | 46 | 6 | 40 | 13.0 |
| 19 | + | M | $v^1$ $xdh1^{\Delta 2}$ | 50 | 0 | 50 | 0 |
| 20 | + | M | $v^1$ $xdh1^{\Delta 2}$ | sterile | 0 | 0 | NA |

| mRNA | donor plasmid |
|---|---|

| 1xNLS-hyPBase | 9xP3-mScarlet-I-SV40T | | | | | | |
|---|---|---|---|---|---|---|---|
| G0 indiv# | mScarlet-I expression | sex | crossed with | Screened G1 animals | RFP**+** | RFP**-** | GTR (%) |
| 1 | + | M | $v^{1}$ $xdh1^{\Delta2}$ | 51 | 0 | 51 | 0 |
| 2 | + | F | $v^{1}$ $xdh1^{\Delta2}$ | 55 | 0 | 55 | 0 |
| 3 | + | F | $v^{1}$ $xdh1^{\Delta2}$ | 60 | 60 | 0 | 100.0 |
| 4 | + | F | $v^{1}$ $xdh1^{\Delta2}$ | 56 | 0 | 56 | 0 |
| 5 | + | F | $v^{1}$ $xdh1^{\Delta2}$ | 52 | 0 | 52 | 0 |
| 6 | + | M | $v^{1}$ $xdh1^{\Delta2}$ | 50 | 0 | 50 | 0 |
| 7 | + | F | $v^{1}$ $xdh1^{\Delta2}$ | sterile | 0 | 0 | NA |
| 8 | + | F | $v^{1}$ $xdh1^{\Delta2}$ | 50 | 50 | 0 | 100.0 |
| 9 | + | M | $v^{1}$ $xdh1^{\Delta2}$ | 53 | 53 | 0 | 100.0 |
| 10 | + | M | $v^{1}$ $xdh1^{\Delta2}$ | 51 | 0 | 51 | 0 |

**Table S4. Results of crossing experiments with 3×NLS-hyPBase.** F, female. M, male. GTR, germ line transformation rate. Data from two independent injection experiments are shown.

| mRNA | donor plasmid | | | | | | |
|---|---|---|---|---|---|---|---|
| 3xNLS-hyPBase | 9xP3-mScarlet-I-SV40T | | | | | | |
| **G0 indiv#** | **mScarlet-I expression** | **sex** | **crossed with** | **Screened G1 animals** | **RFP+** | **RFP-** | **GTR (%)** |
| 1 | + | M | $v^1$ $xdh1^{\Delta 2}$ | 60 | 0 | 60 | 0 |
| 2 | + | M | $v^1$ $xdh1^{\Delta 2}$ | 55 | 0 | 55 | 0 |
| 3 | + | M | $v^1$ $xdh1^{\Delta 2}$ | 3 | 3 | 0 | 100.0 |
| 4 | + | F | $v^1$ $xdh1^{\Delta 2}$ | 51 | 33 | 18 | 64.7 |
| 18 | + | M | $v^1$ $xdh1^{\Delta 2}$ | 61 | 0 | 61 | 0 |
| 19 | + | M | $v^1$ $xdh1^{\Delta 2}$ | 1 | 0 | 1 | 0 |
| 20 | + | M | $v^1$ $xdh1^{\Delta 2}$ | 57 | 0 | 57 | 0 |
| 41 | + | M | $v^1$ $xdh1^{\Delta 2}$ | 30 | 0 | 30 | 0 |

| mRNA | donor plasmid |
|---|---|
| 3xNLS-hyPBase | 9xP3-mScarlet-I-SV40T |

| G0 indiv# | mScarlet-I expression | sex | crossed with | Screened G1 animals | RFP+ | RFP- | GTR (%) |
|---|---|---|---|---|---|---|---|
| 18 | + | M | $v^1$ $xdh1^{\Delta2}$ | 51 | 0 | 51 | 0 |
| 19 | + | M | $v^1$ $xdh1^{\Delta2}$ | sterile | 0 | 0 | NA |
| 20 | + | M | $v^1$ $xdh1^{\Delta2}$ | 57 | 0 | 57 | 0 |
| 21 | + | M | $v^1$ $xdh1^{\Delta2}$ | 68 | 0 | 68 | 0 |
| 22 | + | M | $v^1$ $xdh1^{\Delta2}$ | 64 | 64 | 0 | 100.0 |
| 23 | + | M | $v^1$ $xdh1^{\Delta2}$ | 36 | 0 | 36 | 0 |
| 24 | + | F | $v^1$ $xdh1^{\Delta2}$ | 34 | 0 | 34 | 0 |
| 25 | + | F | $v^1$ $xdh1^{\Delta2}$ | 23 | 0 | 23 | 0 |
| 26 | + | F | $v^1$ $xdh1^{\Delta2}$ | died | 0 | 0 | NA |
| 27 | + | M | $v^1$ $xdh1^{\Delta2}$ | 45 | 0 | 45 | 0 |
| 28 | + | M | $v^1$ $xdh1^{\Delta2}$ | sterile | 0 | 0 | NA |
| 29 | + | F | $v^1$ $xdh1^{\Delta2}$ | 50 | 50 | 0 | 100.0 |
| 30 | + | F | $v^1$ $xdh1^{\Delta2}$ | 54 | 23 | 31 | 42.6 |
| 31 | + | F | $v^1$ $xdh1^{\Delta2}$ | 51 | 51 | 0 | 100.0 |
| 40 | + | F | $v^1$ $xdh1^{\Delta2}$ | 11 | 11 | 0 | 100.0 |

**Table S5. Results of crossing experiments for attP strains.** The integration site was determined for the line derived from G1 individual 8A. F, female. M, male. GTR, germ line transformation rate. Data from two independent injection experiments are shown.

| mRNA | donor plasmid | | | | | | |
|---|---|---|---|---|---|---|---|
| 3xNLS-hyPBase | 9xP3-EGFP-SV40T;attP | | | | | | |
| G0 indiv# | EGFP expression | sex | crossed with | Screened G1 animals | GFP**+** | GFP**-** | GTR (%) |
| 1 | + | F | $v^{1}$ $xdh1^{\Delta2}$ | 49 | 0 | 49 | 0 |
| 2 | + | F | $v^{1}$ $xdh1^{\Delta2}$ | 41 | 0 | 41 | 0 |
| 3 | + | F | $v^{1}$ $xdh1^{\Delta2}$ | died | 0 | 0 | NA |
| 4 | + | F | $v^{1}$ $xdh1^{\Delta2}$ | died | 0 | 0 | NA |
| 5 | + | F | $v^{1}$ $xdh1^{\Delta2}$ | 25 | 19 | 6 | 76.0 |
| 6 | + | M | $v^{1}$ $xdh1^{\Delta2}$ | 26 | 0 | 26 | 0 |
| 7 | + | M | $v^{1}$ $xdh1^{\Delta2}$ | 44 | 0 | 44 | 0 |
| 8 | + | F | $v^{1}$ $xdh1^{\Delta2}$ | 42 | 16 | 26 | 38.1 |
| 9 | + | M | $v^{1}$ $xdh1^{\Delta2}$ | sterile | 0 | 0 | NA |
| 10 | + | M | $v^{1}$ $xdh1^{\Delta2}$ | 5 | 5 | 0 | 100.0 |
| G1 indiv# | EGFP expression | sex | crossed with | Screened G2 animals | GFP**+** | GFP**-** | G2 inheritance rate (%) |
| 5A | + | F | $v^{1}$ $xdh1^{\Delta2}$ | 33 | 27 | 6 | 81.8 |
| 8A | + | F | $v^{1}$ $xdh1^{\Delta2}$ | 64 | 32 | 32 | 50.0 |

| mRNA | donor plasmid | | | | | | |
|---|---|---|---|---|---|---|---|
| 3xNLS-hyPBase | 9xP3-EGFP-SV40T;attP | | | | | | |
| G0 indiv# | EGFP expression | sex | crossed with | Screened G1 animals | GFP**+** | GFP**-** | GTR (%) |
| 1 | + | F | $v^{1}$ $xdh1^{\Delta 2}$ | 48 | 0 | 48 | 0 |
| 2 | + | M | $v^{1}$ $xdh1^{\Delta 2}$ | 31 | 0 | 31 | 0 |
| 3 | + | F | $v^{1}$ $xdh1^{\Delta 2}$ | 45 | 31 | 14 | 68.9 |
| 4 | + | M | $v^{1}$ $xdh1^{\Delta 2}$ | 39 | 0 | 39 | 0 |
| 5 | + | M | $v^{1}$ $xdh1^{\Delta 2}$ | 33 | 6 | 27 | 18.2 |
| G1 indiv# | EGFP expression | sex | crossed with | Screened G2 animals | GFP**+** | GF**P-** | G2 inheritance rate (%) |
| 3A | + | M | $v^{1}$ $xdh1^{\Delta 2}$ | 44 | 37 | 7 | 84.1 |
| 3B | + | F | $v^{1}$ $xdh1^{\Delta 2}$ | 51 | 37 | 14 | 72.5 |

**Table S6. Results of crossing experiments for Gbact-H2B-mScarlet3 strain.** F, female. GTR, germ line transformation rate.

| mRNA | donor plasmid | | | | | | |
|---|---|---|---|---|---|---|---|
| 1xNLS-hyPBase | Gbact-H2B-mScarlet3-SV40T | | | | | | |
| G0 indiv# | mScarlet-I expression | sex | crossed with | Screened G1 animals | RFP**+** | RF**P-** | GTR (%) |
| 1 | + | F | *$v^{1}$ $xdh1^{\Delta 2}$* | 30 | 14 | 16 | 46.7% |

**Table S7. Results of crossing experiments for Gbact-GAP43-mScarlet3;9xP3-EGFP strains.** F, female. M, male. GTR, germ line transformation rate.

| mRNA | donor plasmid | | | | | | |
|---|---|---|---|---|---|---|---|
| 1xNLS-hyPBase | Gbact-GAP43-mScarlet3-SV40T;9xP3-EGFP-SV40T | | | | | | |
| G0 indiv# | EGFP expression | sex | crossed with | Screened G1 animals | GFP**+** | GFP**-** | GTR (%) |
| 1 | + | M | $v^{1}$ $xdh1^{\Delta 2}$ | 45 | 27 | 18 | 60.0 |
| 2 | + | M | $v^{1}$ $xdh1^{\Delta 2}$ | died | 0 | 0 | NA |
| 3 | + | M | $v^{1}$ $xdh1^{\Delta 2}$ | died | 0 | 0 | NA |
| 4 | + | M | $v^{1}$ $xdh1^{\Delta 2}$ | died | 0 | 0 | NA |
| 5 | + | M | $v^{1}$ $xdh1^{\Delta 2}$ | 35 | 0 | 35 | |
| 6 | + | M | $v^{1}$ $xdh1^{\Delta 2}$ | 36 | 0 | 36 | |
| 7 | + | M | $v^{1}$ $xdh1^{\Delta 2}$ | 33 | 0 | 33 | |
| 8 | + | F | $v^{1}$ $xdh1^{\Delta 2}$ | 42 | 23 | 19 | 54.8 |
| 9 | + | F | $v^{1}$ $xdh1^{\Delta 2}$ | 35 | 0 | 35 | |
| 10 | + | F | $v^{1}$ $xdh1^{\Delta 2}$ | died | 0 | 0 | NA |
| 11 | + | F | $v^{1}$ $xdh1^{\Delta 2}$ | 44 | 19 | 25 | 43.2 |
| G1 indiv# | EGFP expression | sex | crossing with | Screened G2 animals | GFP**+** | GFP**-** | G2 inheritance rate (%) |
| 1A | + | M | $v^{1}$ $xdh1^{\Delta 2}$ | 56 | 28 | 28 | 50.0 |
| 1B | + | M | $v^{1}$ $xdh1^{\Delta 2}$ | 46 | 21 | 25 | 45.7 |
| 8A | + | M | $v^{1}$ $xdh1^{\Delta 2}$ | 45 | 23 | 22 | 51.1 |

**Table S8. Results of crossing experiments for Ofact-H2B-mScarlet3 strain.** F, female. M, male. GTR, germ line transformation rate. Data from three independent injection experiments are shown.

| mRNA | donor plasmid | | | | | | |
|---|---|---|---|---|---|---|---|
| 3xNLS-hyPBase | Ofact-H2B-mScarlet3-SV40T | | | | | | |
| G0 indiv# | RFP expression | sex | crossed with | Screened G1 animals | RFP+ | RFP- | GTR (%) |
| 1 | + | F | $v^{1}$ $xdh1^{\Delta 2}$ | 20 | 0 | 20 | 0 |
| 2 | + | F | $v^{1}$ $xdh1^{\Delta 2}$ | 32 | 17 | 15 | 53.1 |
| 3 | + | F | $v^{1}$ $xdh1^{\Delta 2}$ | 20 | 0 | 20 | 0 |
| 4 | + | F | $v^{1}$ $xdh1^{\Delta 2}$ | 49 | 0 | 49 | 0 |
| G1 indiv# | RFP expression | sex | crossed with | Screened G2 animals | RFP+ | RFP- | G2 inheritance rate (%) |
| 1A | + | F | $v^{1}$ $xdh1^{\Delta 2}$ | 50 | 32 | 18 | 64.0 |
| 1B | + | M | $v^{1}$ $xdh1^{\Delta 2}$ | sterile | 0 | 0 | NA |
| 1C | + | M | $v^{1}$ $xdh1^{\Delta 2}$ | sterile | 0 | 0 | NA |
| 1D | + | M | $v^{1}$ $xdh1^{\Delta 2}$ | 31 | 14 | 17 | 45.2 |
| 1E | + | F | $v^{1}$ $xdh1^{\Delta 2}$ | sterile | 0 | 0 | NA |
| 1F | + | F | $v^{1}$ $xdh1^{\Delta 2}$ | 46 | 24 | 22 | 52.2 |

| mRNA | donor plasmid | | | | | | |
|---|---|---|---|---|---|---|---|
| 3xNLS-hyPBase | Ofact-H2B-mScarlet3-SV40T | | | | | | |
| G0 indiv# | RFP expression | sex | crossed with | Screened G1 animals | RFP+ | RFP- | GTR (%) |
| 1 | + | F | $v^{1}\ xdh1^{\Delta 2}$ | 30 | 0 | 30 | 0.0 |
| 2 | + | M | $v^{1}\ xdh1^{\Delta 2}$ | 51 | 10 | 41 | 19.6 |
| G1 indiv# | RFP expression | sex | crossed with | Screened G2 animals | RFP+ | RFP- | G2 inheritance rate (%) |
| 1A | + | F | $v^{1}\ xdh1^{\Delta 2}$ | 53 | 26 | 27 | 49.1 |
| 1B | + | M | $v^{1}\ xdh1^{\Delta 2}$ | sterile | 0 | 0 | NA |
| 1C | + | M | $v^{1}\ xdh1^{\Delta 2}$ | 55 | 33 | 22 | 60.0 |

| mRNA | donor plasmid | | | | | | |
|---|---|---|---|---|---|---|---|
| 3xNLS-hyPBase | Ofact-H2B-mScarlet3-SV40T | | | | | | |
| G0 indiv# | RFP expression | sex | crossed with | Screened G1 animals | RFP+ | RFP- | GTR (%) |
| 1 | + | M | $v^{1}\ xdh1^{\Delta 2}$ | died | 0 | 0 | NA |
| 2 | + | M | $v^{1}\ xdh1^{\Delta 2}$ | died | 0 | 0 | NA |
| 3 | + | M | $v^{1}\ xdh1^{\Delta 2}$ | no data | 3 | no data | no data |
| G1 indiv# | RFP expression | sex | crossed with | Screened G2 animals | RFP+ | RFP- | G2 inheritance rate (%) |
| 3A, 3B, 3C | + | F (3) | $v^{1}\ xdh1^{\Delta 2}$ | 63 | 32 | 31 | 50.8 |

**Table S9. Primers used in the study.**

| Name | Sequence (5'- to -3') | Purpose |
|---|---|---|
| Of-xdh1-F1 | GGCTAACTTCCTAATGGTGGGTT | Genotyping of *Of-xdh1* |
| Of-xdh1-R1 | CTGCCATAGCAGAAAGAAAGGT | Genotyping of *Of-xdh1* |
| hyPBase-Nterm3NLS-F | CCAAAAAAAAGCGCAAGGTTCCGAGATCTCCTGAATTCGC | inverse PCR for pCMV-3xNLS-hyPBase construction |
| hyPBase-Nterm3NLS-R | ACTTTGCGTTTCTTCTTCGGCATGGTGGATCCCATGGTGG | inverse PCR for pCMV-3xNLS-hyPBase construction |
| 3NLS-top | CCGAAGAAGAAACGCAAAGTTCCGAAAAAGAAACGCAAAGTGCCCAAAAAAAAGCGCAAG | Oligonucleotides annealing for pCMV-3xNLS-hyPBase construction |
| 3NLS-bottom | CTTGCGCTTTTTTTTGGGCACTTTGCGTTTCTTTTTCGGAACTTTGCGTTTCTTCTTCGG | Oligonucleotides annealing for pCMV-3xNLS-hyPBase construction |
| SDM-1NLS-F | CCGAGATCTCCTGAATTCGC | Site-directed mutagenesis for pCMV-1xNLS-hyPBase construction |
| SDM-1NLS-R | ACTTTGCGTTTCTTCTTCGG | Site-directed mutagenesis for pCMV-1xNLS-hyPBase construction |
| SDM-5NLS-F | CCTAAGAAAAAGCGTAAAGTCCCGAGATCTCCTGAATTCGC | Site-directed mutagenesis for pCMV-5xNLS-hyPBase construction |

| | | |
|---|---|---|
| SDM-5NLS-R | TACCTTTCGCTTCTTTTTTGGAACCTTGCGCTTTTTTTTTGG | Site-directed mutagenesis for pCMV-5xNLS-hyPBase construction |
| SPLNK-BOT | CGAAGAGTAACCGTTGCTAGGAGAGACCGTGGCTGAATGAGACTGGTGTCGACACTAGTGG | Annealing of splinkerrette PCR |
| SPLNK-GATC-TOP | GATCCCACTAGTGTCGACACCAGTCTCTAATTTTTTTTTTCAAAAAAA | Annealing of splinbkerrette PCR |
| SPLNK#1 | CGAAGAGTAACCGTTGCTAGGAGAGACC | 1st PCR of splinbkerrette PCR |
| SPLNK#2 | GTGGCTGAATGAGACTGGTGTCGAC | 2nd PCR of splinbkerrette PCR |
| 5'SPLNK-PB#1 | ACCGCATTGACAAGCACG | 1st PCR of splinbkerrette PCR |
| 5'SPLNK-PB#2 | CTCCAAGCGGCGACTGAG | 2nd PCR of splinbkerrette PCR |
| 5'SPLNK-PB-SEQ | CGACTGAGATGTCCTAAATGC | Sequencing for splinbkerrette PCR |

**Data S1. plasmid constructs_260728_V4**

pXL[9xP3-mScarlet-I-SV40T] = 1775 bp

TTAACCCTAGAAAGATAATCATATTGTGACGTACGTTAAAGATAATCATGTGTAAAATTGACGC
ATGTGTTTTATCGGTCTGTATATCGAGGTTTATTTATTAATTTGAATAGATATTAAGTTTTATT
ATATTTACACTTACATACTAATAATAAATTCAACAAACAATTTATTTATGTTTATTTATTTATT
AAAAAAAACAAAAACTCAAAATTTCTTCTATAAAGTAACAAAACTTTTATCGAATTCCTGCAGC
CCGGGGGATCCACTAGTTCTGTGAATTCAGATCTCCGGGGATCTAATTCAATTAGAGACTAATT
CAATTAGAGCTAATTCAATTAGGATTAATTCAATTAGAGACTAATTCAATTAGAGCTAATTCAA
TTAGGATTAATTCAATTAGAGACTAATTCAATTAGAGCTAATTCAATTAGGATCCAAGCTTATC
GATTTCGAACCCTCGACCGCCGGAGTATAAATAGAGGCGCTTCGTCTACGGAGCGACAATTCAA
TTCAAACAAGCAAAGTGAACACGTCGCTAAGCGAAAGCTAAGCAAATAAACAAGCGCAGCTGAA
CAAGCTAAACAATCGAAGAATCAAAATGGTGTCTAAGGGAGAGGCTGTCATCAAGGAATTCATG
CGCTTCAAGGTGCACATGGAGGGATCAATGAACGGTCACGAGTTCGAAATCGAGGGAGAAGGAG
AGGGCCGTCCTTACGAAGGTACCCAGACTGCCAAGCTGAAGGTCACCAAGGGTGGCCCACTGCC
TTTCTCCTGGGACATCCTGAGCCCCCAGTTCATGTACGGATCCCGCGCTTTCACTAAGCACCCC
GCCGACATCCCAGACTACTACAAGCAGAGCTTCCCAGAGGGTTTCAAGTGGGAACGTGTGATGA
ACTTCGAGGACGGAGGTGCTGTGACCGTCACTCAGGACACCTCCCTGGAAGACGGCACTCTGAT
CTACAAGGTGAAGCTGAGGGGTACCAACTTCCCTCCCGACGGCCCAGTCATGCAGAAGAAGACC
ATGGGATGGGAGGCTAGCACTGAAAGACTGTACCCTGAGGACGGCGTCCTGAAGGGAGACATCA
AGATGGCCCTGAGGCTGAAGGACGGCGGAAGATACCTGGCTGACTTCAAGACCACTTACAAGGC
CAAGAAGCCTGTGCAGATGCCCGGTGCCTACAACGTCGACAGGAAGCTGGACATCACCTCCCAC
AACGAGGACTACACTGTGGTCGAACAGTACGAGCGCTCTGAAGGCCGTCACTCAACTGGTGGCA
TGGACGAACTGTACAAGTAAAACTTGTTTATTGCAGCTTATAATGGTTACAAATAAAGCAATAG
CATCACAAATTTCACAAATAAAGCATTTTTTTCACTGCATTCTAGTTGTGGTTTGTCCAAACTC
ATCAATGTATCTTAAGATCTTCTAGATATCGGATCCCGGGGGTCGACGGTATCGATAAGCTTGA

TATCTATAACAAGAAAATATATATATAATAAGTTATCACGTAAGTAGAACATGAAATAACAATA
TAATTATCGTATGAGTTAAATCTTAAAAGTCACGTAAAAGATAATCATGCGTCATTTTGACTCA
CGCGGTCGTTATAGTTCAAAATCAGTGACACTTACCGCATTGACAAGCACGCCTCACGGGAGCT
CCAAGCGGCGACTGAGATGTCCTAAATGCACAGCGACGGATTCGCGCTATTTAGAAAGAGAGAG
CAATATTTCAAGAATGCATGCGTCAATTTTACGCAGACTATCTTTCTAGGGTTAA

*piggybac* ITR: 237* bp(L), 306 bp(R)

* 1 adenine deletion

Pax-6 homeodomain binding site: 11 bp x 9

*Drosophila hsp70* promoter (minimal): 125 bp

mScarlet-I: 696 bp

SV40 polyadenylation signal: 122 bp

pXL[9xP3-EGFP-SV40T;attP] = 1838 bp

```
TTAACCCTAGAAAGATAATCATATTGTGACGTACGTTAAAGATAATCATGTGTAAAATTGACGC
ATGTGTTTTATCGGTCTGTATATCGAGGTTTATTTATTAATTTGAATAGATATTAAGTTTTATT
ATATTTACACTTACATACTAATAATAAATTCAACAAACAATTTATTTATGTTTATTTATTTATT
AAAAAAAACAAAAACTCAAAATTTCTTCTATAAAGTAACAAAACTTTTATCGAATTCCTGCAGC
CCGGGGGATCCACTAGTTCTGTGAATTCAGATCTCCGGGGATCTAATTCAATTAGAGACTAATT
CAATTAGAGCTAATTCAATTAGGATTAATTCAATTAGAGACTAATTCAATTAGAGCTAATTCAA
TTAGGATTAATTCAATTAGAGACTAATTCAATTAGAGCTAATTCAATTAGGATCCAAGCTTATC
GATTTCGAACCCTCGACCGCCGGAGTATAAATAGAGGCGCTTCGTCTACGGAGCGACAATTCAA
TTCAAACAAGCAAAGTGAACACGTCGCTAAGCGAAAGCTAAGCAAATAAACAAGCGCAGCTGAA
CAAGCTAAACAATCGAAGAATCAAAATGGTGAGCAAGGGCGAGGAGCTGTTCACCGGGGTGGTG
CCCATCCTGGTCGAGCTGGACGGCGACGTAAACGGCCACAAGTTCAGCGTGTCCGGCGAGGGCG
AGGGCGATGCCACCTACGGCAAGCTGACCCTGAAGTTCATCTGCACCACCGGCAAGCTGCCCGT
GCCCTGGCCCACCCTCGTGACCACCCTGACCTACGGCGTGCAGTGCTTCAGCCGCTACCCCGAC
CACATGAAGCAGCACGACTTCTTCAAGTCCGCCATGCCCGAAGGCTACGTCCAGGAGCGCACCA
TCTTCTTCAAGGACGACGGCAACTACAAGACCCGCGCCGAGGTGAAGTTCGAGGGCGACACCCT
GGTGAACCGCATCGAGCTGAAGGGCATCGACTTCAAGGAGGACGGCAACATCCTGGGGCACAAG
CTGGAGTACAACTACAACAGCCACAACGTCTATATCATGGCCGACAAGCAGAAGAACGGCATCA
AGGTGAACTTCAAGATCCGCCACAACATCGAGGACGGCAGCGTGCAGCTCGCCGACCACTACCA
GCAGAACACCCCCATCGGCGACGGCCCCGTGCTGCTGCCCGACAACCACTACCTGAGCACCCAG
TCCGCCCTGAGCAAAGACCCCAACGAGAAGCGCGATCACATGGTCCTGCTGGAGTTCGTGACCG
CCGCCGGGATCACTCTCGGCATGGACGAGCTGTACAAGTAAAACTTGTTTATTGCAGCTTATAA
TGGTTACAAATAAAGCAATAGCATCACAAATTTCACAAATAAAGCATTTTTTTCACTGCATTCT
AGTTGTGGTTTGTCCAAACTCATCAATGTATCTTAAGATCTTCTAGATATCGGATCCCGGTGCC
CCAACTGGGGTAACCTTTGAGTTCTCTCAGTTGGGGGGGGGTCGACGGTATCGATAAGCTTGAT
```

ATCTATAACAAGAAAATATATATATAATAAGTTATCACGTAAGTAGAACATGAAATAACAATAT
AATTATCGTATGAGTTAAATCTTAAAAGTCACGTAAAAGATAATCATGCGTCATTTTGACTCAC
GCGGTCGTTATAGTTCAAAATCAGTGACACTTACCGCATTGACAAGCACGCCTCACGGGAGCTC
CAAGCGGCGACTGAGATGTCCTAAATGCACAGCGACGGATTCGCGCTATTTAGAAAGAGAGAGC
AATATTTCAAGAATGCATGCGTCAATTTTACGCAGACTATCTTTCTAGGGTTAA

*piggybac* ITR: 237* bp(L), 305 bp(R)
* 1 adenine deletion

Pax-6 homeodomain binding site: 11 bp x 9

*Drosophila hsp70* promoter (minimal): 125 bp

EGFP: 717 bp

SV40 polyadenylation signal: 122 bp

phage φC31 attP: 42 bp

pXL[Gbact-H2B-mScarlet3-SV40T] = 2974 bp

TTAACCCTAGAAAGATAATCATATTGTGACGTACGTTAAAGATAATCATGTGTAAAATTGACGC

ATGTGTTTTATCGGTCTGTATATCGAGGTTTATTTATTAATTTGAATAGATATTAAGTTTTATT

ATATTTACACTTACATACTAATAATAAATTCAACAAACAATTTATTTATGTTTATTTATTTATT

AAAAAAAAACAAAAACTCAAAATTTCTTCTATAAAGTAACAAAACTTTTATCGAATTCCTGCAG

CCCGGGGGATCCACTAGCGCTACCGGACTCAGATCTCGAGCAGTTCCTTCAAATGTTCAAAATA

GTATTAGTTTTTGTTACGCCTAAATTTCTTGAGAATGTTGTTTAAACTTTTATTTTCTGACAAA

TGTTAAAATCATTGCTGTTTTTTTAGCAGACACTTTTCGAAGCTTGTGTTTTGATATCTCGTTG

TAAAGTAAATTCATTTATAAGTTCTCATTGACTTTTAACCGTTTTTAGCCGGCATGTTCCGGTA

GGGAGCGTTGGTTCCACGCTGCTGGCTCGATACCATATATGGTAAAGAAAGCTGCCATTGGATG

AGAAGCATGCCCTTATTAGTTGGTGGTGGGGGTAGGGTAGGCATGGGAGGGGCAGGGCCTCCGT

AGTATATTACTCATAGGCCGGCTCCCGGCTTGAATCAGTCTGGTGCTAGTTCTCGTACTGTGAA

GATAGCAGTGTTGCGCTGCTGCGTTTTTGTACATATTACATTGTAAGTTTCTAGTGCTTCTCGA

AGATACTTTGTGAATTTGTGTTACGAGCATTTGTGCCATGTGGGATATCAGCACAATTTTTCGT

GTGGATTTTCGCGTATGTGCTTGTGAGCAGTTAGGGCGGGGCAACCGGCTTCGTCCACACTCCC

AAGGATTTCGCCTTATTTGGCGGGTGTTTGCTTAGGGTCTAGCCACTGAATATGGGCGGTGCCT

GTGTAGTGGTTGCCGGTGTGGGGCTTAGTCTTGTCAATGGGCGGGAAACCGGTGTGGGGAAACT

AGTAGCGAATTGGATGGTTAACTAGTTTACTCGTGCTACGGTTCTCTTAATTTTTTTTTTATTT

CTTGCAGTACCCGTAAACTCAACTACTAACCGAATTCTGCAGGTCGACATGCCGCCCAAGACTA

GCGGAAAGGCCGCGAAGAAGGCCGGTAAGGCCCAGAAGAATATCTCCAAGGGAGACAAGAAGAA

GAAGCGCAAGAGGAAGGAAAGCTACGCCATCTACATCTACAAGGTGTTGAAGCAGGTCCATCCT

GACACCGGCATCTCCAGCAAGGCGATGAGCATCATGAACAGTTTTGTCAATGACATTTTCGAGC

GCATCGCCGCCGAGGCTTCCCGCCTGGCGCACTACAACAAGCGCTCTACCATCACCTCGCGGGA

GATCCAGACCGCTGTGCGTCTCCTTCTGCCCGGCGAGCTGGCGAAGCACGCTGTCAGCGAAGGC

ACTAAGGCTGTCACCAAGTACACCAGCTCCAAGAAGGATCCACCGGTCGCCACCATGGATAGCA

CCGAGGCAGTGATCAAGGAGTTCATGCGGTTCAAGGTGCACATGGAGGGCTCCATGAACGGCCA
CGAGTTCGAGATCGAGGGCGAGGGCGAGGGCCGCCCCTACGAGGGCACCCAGACCGCCAAGCTG
AGGGTGACCAAGGGTGGCCCCCTGCCCTTCTCCTGGGACATCCTGTCCCCTCAGTTCATGTACG
GCTCCAGGGCCTTCACGAAGCACCCCGCCGACATCCCCGACTACTGGAAGCAGTCCTTCCCCGA
GGGCTTCAAGTGGGAGCGCGTGATGAACTTCGAGGACGGCGGCGCTGTGTCCGTGGCCCAGGAC
ACCTCCCTGGAGGACGGCACCCTGATCTACAAGGTGAAGCTCCGCGGCACCAACTTCCCTCCTG
ACGGCCCCGTAATGCAGAAGAAGACAATGGGCTGGGAAGCATCCACCGAGCGGTTGTACCCCGA
GGACGTCGTGCTGAAGGGCGACATTAAGATGGCCCTGCGCCTGAAGGACGGCGGTCGCTACCTG
GCGGACTTCAAGACCACCTACAGGGCCAAGAAGCCCGTGCAGATGCCCGGCGCTTTCAACATCG
ACCGCAAGTTGGACATCACATCCCACAACGAGGACTACACCGTGGTGGAACAGTACGAACGCTC
CGTGGCCCGCCACTCCACCGGCTAAAGCGGCCGCGACTCTAGATCATAATCAGCCATACCACAT
TTGTAGAGGTTTTACTTGCTTTAAAAAACCTCCCACACCTCCCCCTGAACCTGAAACATAAAAT
GAATGCAATTGTTGTTGTTAACTTGTTTATTGCAGCTTATAATGGTTACAAATAAAGCAATAGC
ATCACAAATTTCACAAATAAAGCATTTTTTTCACTGCATTCTAGTTGTGGTTTGTCCAAACTCA
TCAATGTATCTTAAGGCGTAAATTGTAAGCGTTAATATTTTGTTAAAATTCGCGTTAAATTTTT
GTTAAATCAGCTCATTTTTTAACCAATAGGCCGAAATCGGCAAAATCCCTTATAAATCAAAAGA
ATAGACCGAGATAGGGTTGAGTGTTGTTCCAGTTAGATCTTAATACGACTCACTATAGGGCGAA
TTGGGTACCGGGCCCCCCCTCGAGGTCGACGGTATCGATAAGCTTGATATCTATAACAAGAAAA
TATATATATAATAAGTTATCACGTAAGTAGAACATGAAATAACAATATAATTATCGTATGAGTT
AAATCTTAAAAGTCACGTAAAAGATAATCATGCGTCATTTTGACTCACGCGGTCGTTATAGTTC
AAAATCAGTGACACTTACCGCATTGACAAGCACGCCTCACGGGAGCTCCAAGCGGCGACTGAGA
TGTCCTAAATGCACAGCGACGGATTCGCGCTATTTAGAAAGAGAGAGCAATATTTCAAGAATGC
ATGCGTCAATTTTACGCAGACTATCTTTCTAGGGTTAA

*piggybac* ITR: 238 bp(L), 305 bp(R)

*Gryllus actin* promoter: 826 bp

*Gryllus* H2B: 369 bp

mScarlet3: 672 bp

SV40 polyadenylation signal: 122 bp

pXL[Gbact-gap43-mScarlet3-SV40T;9xP3-EGFP-SV40T] = 3635 bp

```
TTAACCCTAGAAAGATAATCATATTGTGACGTACGTTAAAGATAATCATGTGTAAAATTGACGC
ATGTGTTTTATCGGTCTGTATATCGAGGTTTATTTATTAATTTGAATAGATATTAAGTTTTATT
ATATTTACACTTACATACTAATAATAAATTCAACAAACAATTTATTTATGTTTATTTATTTATT
AAAAAAAAACAAAAACTCAAAATTTCTTCTATAAAGTAACAAAACTTTTATCGAATTCCTGCAG
CCCGGGGGATCCACTAGCGCTACCGGACTCAGATCTCGAGCAGTTCCTTCAAATGTTCAAAATA
GTATTAGTTTTTGTTACGCCTAAATTTCTTGAGAATGTTGTTTAAACTTTTATTTTCTGACAAA
TGTTAAAATCATTGCTGTTTTTTTAGCAGACACTTTTCGAAGCTTGTGTTTTGATATCTCGTTG
TAAAGTAAATTCATTTATAAGTTCTCATTGACTTTTAACCGTTTTTAGCCGGCATGTTCCGGTA
GGGAGCGTTGGTTCCACGCTGCTGGCTCGATACCATATATGGTAAAGAAAGCTGCCATTGGATG
AGAAGCATGCCCTTATTAGTTGGTGGTGGGGGTAGGGTAGGCATGGGAGGGGCAGGGCCTCCGT
AGTATATTACTCATAGGCCGGCTCCCGGCTTGAATCAGTCTGGTGCTAGTTCTCGTACTGTGAA
GATAGCAGTGTTGCGCTGCTGCGTTTTTGTACATATTACATTGTAAGTTTCTAGTGCTTCTCGA
AGATACTTTGTGAATTTGTGTTACGAGCATTTGTGCCATGTGGGATATCAGCACAATTTTTCGT
GTGGATTTTCGCGTATGTGCTTGTGAGCAGTTAGGGCGGGGCAACCGGCTTCGTCCACACTCCC
AAGGATTTCGCCTTATTTGGCGGGTGTTTGCTTAGGGTCTAGCCACTGAATATGGGCGGTGCCT
GTGTAGTGGTTGCCGGTGTGGGGCTTAGTCTTGTCAATGGGCGGGAAACCGGTGTGGGGAAACT
AGTAGCGAATTGGATGGTTAACTAGTTTACTCGTGCTACGGTTCTCTTAATTTTTTTTTTATTT
CTTGCAGTACCCGTAAACTCAACTACTAACCGAATTCTGCAGGTCGACATGCTGTGCTGTATGA
GAAGAACCAAACAGGTTGAAAAGAATGATGAGGACCAAAAGATCATGGATAGCACCGAGGCAGT
GATCAAGGAGTTCATGCGGTTCAAGGTGCACATGGAGGGCTCCATGAACGGCCACGAGTTCGAG
ATCGAGGGCGAGGGCGAGGGCCGCCCCTACGAGGGCACCCAGACCGCCAAGCTGAGGGTGACCA
AGGGTGGCCCCCTGCCCTTCTCCTGGGACATCCTGTCCCCTCAGTTCATGTACGGCTCCAGGGC
CTTCACGAAGCACCCCGCCGACATCCCCGACTACTGGAAGCAGTCCTTCCCCGAGGGCTTCAAG
```

TGGGAGCGCGTGATGAACTTCGAGGACGGCGGCGCTGTGTCCGTGGCCCAGGACACCTCCCTGG
AGGACGGCACCCTGATCTACAAGGTGAAGCTCCGCGGCACCAACTTCCCTCCTGACGGCCCCGT
AATGCAGAAGAAGACAATGGGCTGGGAAGCATCCACCGAGCGGTTGTACCCCGAGGACGTCGTG
CTGAAGGGCGACATTAAGATGGCCCTGCGCCTGAAGGACGGCGGTCGCTACCTGGCGGACTTCA
AGACCACCTACAGGGCCAAGAAGCCCGTGCAGATGCCCGGCGCTTTCAACATCGACCGCAAGTT
GGACATCACATCCCACAACGAGGACTACACCGTGGTGGAACAGTACGAACGCTCCGTGGCCCGC
CACTCCACCGGCTAAAGCGGCCGCGACTCTAGATCATAATCAGCCATACCACATTTGTAGAGGT
TTTACTTGCTTTAAAAAACCTCCCACACCTCCCCCTGAACCTGAAACATAAAATGAATGCAATT
GTTGTTGTTAACTTGTTTATTGCAGCTTATAATGGTTACAAATAAAGCAATAGCATCACAAATT
TCACAAATAAAGCATTTTTTTCACTGCATTCTAGTTGTGGTTTGTCCAAACTCATCAATGTATC
TTAAGGTTCTGTGAATTCAGATCTCCGGGGATCTAATTCAATTAGAGACTAATTCAATTAGAGC
TAATTCAATTAGGATTAATTCAATTAGAGACTAATTCAATTAGAGCTAATTCAATTAGGATTAA
TTCAATTAGAGACTAATTCAATTAGAGCTAATTCAATTAGGATCCAAGCTTATCGATTTCGAAC
CCTCGACCGCCGGAGTATAAATAGAGGCGCTTCGTCTACGGAGCGACAATTCAATTCAAACAAG
CAAAGTGAACACGTCGCTAAGCGAAAGCTAAGCAAATAAACAAGCGCAGCTGAACAAGCTAAAC
AATCGAAGAATCAAAATGGTGAGCAAGGGCGAGGAGCTGTTCACCGGGGTGGTGCCCATCCTGG
TCGAGCTGGACGGCGACGTAAACGGCCACAAGTTCAGCGTGTCCGGCGAGGGCGAGGGCGATGC
CACCTACGGCAAGCTGACCCTGAAGTTCATCTGCACCACCGGCAAGCTGCCCGTGCCCTGGCCC
ACCCTCGTGACCACCCTGACCTACGGCGTGCAGTGCTTCAGCCGCTACCCCGACCACATGAAGC
AGCACGACTTCTTCAAGTCCGCCATGCCCGAAGGCTACGTCCAGGAGCGCACCATCTTCTTCAA
GGACGACGGCAACTACAAGACCCGCGCCGAGGTGAAGTTCGAGGGCGACACCCTGGTGAACCGC
ATCGAGCTGAAGGGCATCGACTTCAAGGAGGACGGCAACATCCTGGGGCACAAGCTGGAGTACA
ACTACAACAGCCACAACGTCTATATCATGGCCGACAAGCAGAAGAACGGCATCAAGGTGAACTT
CAAGATCCGCCACAACATCGAGGACGGCAGCGTGCAGCTCGCCGACCACTACCAGCAGAACACC
CCCATCGGCGACGGCCCCGTGCTGCTGCCCGACAACCACTACCTGAGCACCCAGTCCGCCCTGA

GCAAAGACCCCAACGAGAAGCGCGATCACATGGTCCTGCTGGAGTTCGTGACCGCCGCCGGGAT
CACTCTCGGCATGGACGAGCTGTACAAGTAAAACTTGTTTATTGCAGCTTATAATGGTTACAAA
TAAAGCAATAGCATCACAAATTTCACAAATAAAGCATTTTTTTCACTGCATTCTAGTTGTGGTT
TGTCCAAACTCATCAATGTATCTTGGTACCGGGCCCCCCCTCGAGGTCGACGGTATCGATAAGC
TTGATATCTATAACAAGAAAATATATATATAATAAGTTATCACGTAAGTAGAACATGAAATAAC
AATATAATTATCGTATGAGTTAAATCTTAAAAGTCACGTAAAAGATAATCATGCGTCATTTTGA
CTCACGCGGTCGTTATAGTTCAAAATCAGTGACACTTACCGCATTGACAAGCACGCCTCACGGG
AGCTCCAAGCGGCGACTGAGATGTCCTAAATGCACAGCGACGGATTCGCGCTATTTAGAAAGAG
AGAGCAATATTTCAAGAATGCATGCGTCAATTTTACGCAGACTATCTTTCTAGGGTTAA

*piggybac* ITR: 238 bp(L), 305 bp(R)

*Gryllus actin* promoter: 826 bp

GAP43 fragment (palmitoylation sites): 60 bp

mScarlet3: 672 bp

SV40 polyadenylation signal: 122 bp

Pax-6 homeodomain binding site: 11 bp x 9

*Drosophila hsp70* promoter (minimal): 125 bp

EGFP: 717 bp

SV40 polyadenylation signal: 121 bp

pXL[Ofact-H2B-mScarlet3-SV40T] = 4650 bp

TTAACCCTAGAAAGATAATCATATTGTGACGTACGTTAAAGATAATCATGTGTAAAATTGACGC
ATGTGTTTTATCGGTCTGTATATCGAGGTTTATTTATTAATTTGAATAGATATTAAGTTTTATT
ATATTTACACTTACATACTAATAATAAATTCAACAAACAATTTATTTATGTTTATTTATTTATT
AAAAAAAAACAAAAACTCAAAATTTCTTCTATAAAGTAACAAAACTTTTATCGAATTCCTGCAG
CCCGGGGGATCCACTAGCGCTACCGGACTCAGATCTCGAGCGGGGAGATTTCTCGCCTTTCCTT
AGTGAGGAGTGGAGAGAGACTGTGGATATATATATATATATCCACACGTACACATTTATATATG
GTAGATAATATCTCTAAGAATTTGATTCTTTGACTGCACGCCACCGACATGCCAACCATCCAGA
GGTTAATTATATTTAGGAAATTTGGTTTAGGAATCCTTAAAAATAAACATTTGTTAAAACCTCG
ACATAGACTTCCTTCTTGATTCTTATACTTGACACTACATTGTGGTTAATTTAAGGTTGCTCGG
GGAAAATCATTCAGACATGTAATAAAATAGAAACGGATTTTGAACTTAATTTCTGTTTACAGTT
AGATAAAGAATTGTATTGCTATATAATGTACACACTGGACTGTCTGTGCCTGTGCTTATCCTAG
TAAATCGGCTTAACTAATAACCCTTCTTATAGAGGTAGATTGGTTGAGAAGATGACTTTCGTTT
CCAACATTAAAACAGAAACAAAATTACTTCTGTTTGGGTACTTATATAAAACCTGTAAGAAAGC
GAACCATAATAAGCTGTCAAGAAGAGCCCGGACCACAATGCCTAGGTCAGGTAGAAGCTGGTAG
AGAGATTTGGCAGAGGGCAGACCCATTGATGAGAGAATGGGCATGAATGGGAGTTTTAAAGGGA
ATCGGGAGAGTTTTAGAGCCTCAGGGATAGGGACTCTCGCGTCAGACGACATTTCGACAGGGGC
TTAACGGAATTGCACCAGGAATTAGGGATATTGTAGGGGACGGACGGGAGACTTGGAGGGACTC
TCAGCGCTACTAGGATTCGGAAATATTTCCAGTGACAGAACGGGAAATGGACTCCGCCTGCCTG
CAAATGCATGTTCAATAATAAAAACTGGCTGAGAGATATTTACAAAGGCGGCTCTTACTAAATG
CCGGCGAGCTGGAGGGATTTCTCCCTCGCCGCGGCTCCGCCAATCAGCGGGCTCCATCCGTTT
TATGCCCATATATGGTAAGAGCGGCGGTCGAGGCCGCTGGTGGGGGAAGGCGGGTCCACCTCCC
TCCTCCCCCCCGACTCCCGGCTTTTCCCCCTCACAGCTATACCATACTTAACCCAATTGTTGTT
ATAATTTTTTTAATTACTCATTTTTTCCCATAAAAAAAACTGTATCGTGAAATCCGCTACCGGG
AAGGCGTATTATCAGCGGAATAGTGAAGTATTTGTGAGTGGCGGAGGAGGCGACACCCCCCGGA

TCGGGCGAGTCCACTCTCCTCCTGGAGACCAACGGCTCAATCCTCCTTCACTCCCGTCAATTGT
GTGATAGTGTACGCATTGGCTCTCGCCTCCTTCGTCCAGTAAAGGTAAGTCTTGCAGAGAAATC
TCACTATTTCCTACACTTACAGTAAATACTCTACATCCCTACTCTTCTAATAGTACTAACCGTT
TGTATATTGATATTTAAAGTGAAGGGGCCGGACTTTACTCCCGAATGAGTGCCGGTAGGACCTC
GCAAACGTGTTGCCTTCGTTTTCATTCTCTCGTCCTCTAAGTAAATTTAACTCCGGTATTTCGG
GACGTCCACAGCCAGAATTAAACTTGGATCTCTTCCAATATCTATGTTTAGGGAATGGTTTACT
ACGGTAGCGCCATTGACTGGAGCGCTATTTGTGTACTGATGAGAGTAGAGATGGCTCCTGGCCT
ATCTTGCCCTACCCCAAGTGTGGCGGTAAATCAGTTTTACCGGCTCTTTTAGGCCGGAATTGTG
TATTCCCTCGACCGTACTGCCATATAAGGAGATGTGCAGAGAGAGGGCGTGGAGGGGCGGGGGC
CGGTGGAGGGGGACGGCCTATATCACCCTCGGGACCCCTGCCCAAGCCACCAGTCCGACTTCAC
AGCTCAAACAGTACTCTGCGTACTATTTATTTATTTATTTTTAATAAGTAAGTGACTTCTCAAT
AACCTCTTAAAGAGTGGGTGTGGATTATAGTGCTACTTATATTAATCTATTAGTGCTCATCAGT
GAATCGTGTTTTATGTGATATGAGTGATTCAAAGTGAAACTTAATGGATTTACAATTAAATCTT
CAGGGTCAGATTATAGCATTTGGTCAAGCTTCCCGAGAGGTAAAATTTTTCCCTATTCCCTCTA
GGCCAATTGGATTTTAAATCAAGCATCCAATTGTCAAAATTAATTCATCTTATAGCTACTGGAA
ATATCTGGAATTATGTGAAACTCATTCAGTAAAACCCCCCATTTATTTAACATGTGTTATAACT
TAGTTGTATTTTAAATCACATGTACACTGACTTCAATTCACACAGGATGCCTGTAAAGTGATTT
ATTTAGTATTTTTTTTTTACAATATTTAGAATATTTTCAAATTATGTACTGTAAAATGCTTGAC
TATCTGTTGTTCTGAAACCCTTGAAGGCACTCTCTGATTTTATGGATACCTTCCATAGGATTTT
AATTTTAGTTTTTGTTTTTCAGCTTATTTAACTAACCCAAAACAAATTCTGCAGGTCGACATGC
CGCCCAAGACTAGCGGAAAGGCCGCGAAGAAGGCCGGTAAGGCCCAGAAGAATATCTCCAAGGG
AGACAAGAAGAAGAAGCGCAAGAGGAAGGAAAGCTACGCCATCTACATCTACAAGGTGTTGAAG
CAGGTCCATCCTGACACCGGCATCTCCAGCAAGGCGATGAGCATCATGAACAGTTTTGTCAATG
ACATTTTCGAGCGCATCGCCGCCGAGGCTTCCCGCCTGGCGCACTACAACAAGCGCTCTACCAT
CACCTCGCGGGAGATCCAGACCGCTGTGCGTCTCCTTCTGCCCGGCGAGCTGGCGAAGCACGCT

GTCAGCGAAGGCACTAAGGCTGTCACCAAGTACACCAGCTCCAAGAAGGATCCACCGGTCGCCA
CCATGGATAGCACCGAGGCAGTGATCAAGGAGTTCATGCGGTTCAAGGTGCACATGGAGGGCTC
CATGAACGGCCACGAGTTCGAGATCGAGGGCGAGGGCGAGGGCCGCCCCTACGAGGGCACCCAG
ACCGCCAAGCTGAGGGTGACCAAGGGTGGCCCCCTGCCCTTCTCCTGGGACATCCTGTCCCCTC
AGTTCATGTACGGCTCCAGGGCCTTCACGAAGCACCCCGCCGACATCCCCGACTACTGGAAGCA
GTCCTTCCCCGAGGGCTTCAAGTGGGAGCGCGTGATGAACTTCGAGGACGGCGGCGCTGTGTCC
GTGGCCCAGGACACCTCCCTGGAGGACGGCACCCTGATCTACAAGGTGAAGCTCCGCGGCACCA
ACTTCCCTCCTGACGGCCCCGTAATGCAGAAGAAGACAATGGGCTGGGAAGCATCCACCGAGCG
GTTGTACCCCGAGGACGTCGTGCTGAAGGGCGACATTAAGATGGCCCTGCGCCTGAAGGACGGC
GGTCGCTACCTGGCGGACTTCAAGACCACCTACAGGGCCAAGAAGCCCGTGCAGATGCCCGGCG
CTTTCAACATCGACCGCAAGTTGGACATCACATCCCACAACGAGGACTACACCGTGGTGGAACA
GTACGAACGCTCCGTGGCCCGCCACTCCACCGGCTAAAGCGGCCGCGACTCTAGATCATAATCA
GCCATACCACATTTGTAGAGGTTTTACTTGCTTTAAAAAACCTCCCACACCTCCCCCTGAACCT
GAAACATAAAATGAATGCAATTGTTGTTGTTAACTTGTTTATTGCAGCTTATAATGGTTACAAA
TAAAGCAATAGCATCACAAATTTCACAAATAAAGCATTTTTTTCACTGCATTCTAGTTGTGGTT
TGTCCAAACTCATCAATGTATCTTAAGGCGTAAATTGTAAGCGTTAATATTTTGTTAAAATTCG
CGTTAAATTTTTGTTAAATCAGCTCATTTTTTAACCAATAGGCCGAAATCGGCAAAATCCCTTA
TAAATCAAAAGAATAGACCGAGATAGGGTTGAGTGTTGTTCCAGTTAGATCTTAATACGACTCA
CTATAGGGCGAATTGGGTACCGGGCCCCCCCTCGAGGTCGACGGTATCGATAAGCTTGATATCT
ATAACAAGAAAATATATATATAATAAGTTATCACGTAAGTAGAACATGAAATAACAATATAATT
ATCGTATGAGTTAAATCTTAAAAGTCACGTAAAAGATAATCATGCGTCATTTTGACTCACGCGG
TCGTTATAGTTCAAAATCAGTGACACTTACCGCATTGACAAGCACGCCTCACGGGAGCTCCAAG
CGGCGACTGAGATGTCCTAAATGCACAGCGACGGATTCGCGCTATTTAGAAAGAGAGAGCAATA
TTTCAAGAATGCATGCGTCAATTTTACGCAGACTATCTTTCTAGGGTTAA

*piggybac* ITR: 238 bp(L), 305 bp(R)

*Oncopeltus actin* promoter: 2502 bp

*Gryllus* H2B: 369 bp

mScarlet3: 672 bp

SV40 polyadenylation signal: 122 bp

pXL[Ofinv2.4kb-QF2;15xQUAS-mScarlet3-SV40T;9xP3-EGFP-SV40T] = 7161 bp

TTAACCCTAGAAAGATAATCATATTGTGACGTACGTTAAAGATAATCATGTGTAAAATTGACGC
ATGTGTTTTATCGGTCTGTATATCGAGGTTTATTTATTAATTTGAATAGATATTAAGTTTTATT
ATATTTACACTTACATACTAATAATAAATTCAACAAACAATTTATTTATGTTTATTTATTTATT
AAAAAAAAACAAAAACTCAAAATTTCTTCTATAAAGTAACAAAACTTTTATCGAATTCCTGCAG
CCCGGGGGATCCACTAGCGCTACCGGACTCAGATCTCGAGATGATGAGGTGAAGTGTACTCGAA
GTAAGTGGACAGGGCGAGGGTGCTGTGGTCTTGTCAACAATTCTCTTTCTCACACCCACATTTC
ATCAGTTCTTCACTGTGTCTCATAAGATGCTTAATGCTTAAGTATGCTTAGGCCACATCAAAGC
TCTGTAAATAAATGTGCTTTGCGCTTTGTGATAAGGCTTTGCATGACTTAAAATACTAATATAT
ATAAAAAGAAAACTAAATATTCGCAATACTTTGCGTTATTTCTATTTTGTGTGTTTGTGAGTGT
TGACTACTCTCTCATTCTTCCTCCTGTTATTAATATATATATATTTAATATATGACTTTATATA
TTATATACATATAATTTCTAATTTCCGATTGCGTGATTAGACAAAATGTGAATCGCTAGCGTTA
GACAACTATTGCCGACTGCACCACGTATGCCGCCATGTGTATCCTTGGGAAGCGTGTTTCGCTC
CTTGGTCTTCCTAGCTCTTCCTTCCAGTTATCTTTATCTTCAGAGCTTCCTGTCTTATCAAACT
CAACATAATTTCAGCAAGACAACACACACTAAATATTGGACTTTTCCATCTTTCACATTTGCAA
ATCTTCACTACCAATATGTCATACCAATGGCATCATAAATTAACCTGTTGTAAGACTCGATGCC
CCTCTGAAATTGTCAACTAATTTCCTGTGGTTTCCATTTTCCTTGATAGGAATAATACACCTTC
CACAATTAGATAATTATGGTATTATACATATAAACCTACACAGAGCTATCTATTGAAGTACAGA
TTATTTTAAGTTTAGTCTTTTGTGATGGGCTAATCCACGAGGCTGACTGACGTAATTTGGTTTC
ATATCCTACATAATTATCGGACGTGATTTGGGTAGGTCAATATTAGGAAAACCAATATTATATA
TAAATCATTTTTAACTTCTTTTTTTTTTTTTAAGTCGTGGTACAAGCGTGAATATATTGATTT
CTTGTAGGAAATGTACTAAAATCGAATAATCGTATTTATAACAGTTGCCATAAAGTTGAAGTAA
CTGAATAAGTGTAAAATTCAATTCATTACATTTAATGTATAATAACATTTTTAGAGAGTAAGGC

ACTAAAAATTGTAGCTGTGCGGTTGAGCGAACCAACTAACAGGTAAAATTCAAATAAAGCACTT
CATTGCAAAATAGGGGGAGGGGGAGGGGGGAGAGATTTACAGGAGAAACAAATAGAATGCTCGC
AGGAATCGTGGTTATTATGGTAGGTGTATAGGACTAAAATCTTATCTACAAAAGTGCATTATAT
TGTAACAGTTTAATATCCACTGACTTCCAGGACCTCCTTGAGTCATCGGAATGATACAGCTTTA
TTTTATTTCATTTTAGAAAATATTTATGGGAAAAATAAATTGTTTCCAGACAGGATAGCTTGCA
ATGGATTTGAGGATAATAAACTTGTTCTACTGTACTGTGTGTTTCATGATTTCGAAGTTAGGAC
AAAATCAAACTTTGGGATCAGCGCCGCGTACTTGATATAAATTACATTATGTGAAATCTGTTGA
AAACGATTAATTGTATGGTCTTCTAAAGAAGGAAATGATATTTTATAGCCTCCCATCTGAATTG
GAAGGGCCAGTCAACGGAGGTTGCTTTCCATACCAAATTTATATTAATTGCAATTGTCAACTTC
TTAAATTGGACGTAATGAATATTACTTCCCCTACTAAAGGGGTGTTGATGGTAATCGCAACCCT
CGATGAATACGATTGCTCAAAGTGTCTATTGTCTTAGGCTGTTGAGCTCCTAAAAAAGAAAGGC
TGTCAATCGCATTCATTAGCAACGAAAATGATTCTAACTGGATACATAACCACACAGGACGAGA
AAGACCAACTATTTCTTCCTGCGAAAAGAATATTTAGAGGCTAATTTGATTACATTCTTCATTC
AAAGAGAAAACTGATTTCCATCAAAATTAAAAATTAACTGTCAGAACACTCAACGAATTGATAT
GTAATCGATGATACACATCGATGATAGTCTTCCCACATAGATAATTAATTCATGAGTGAGAGGG
CGTGAGTGAGGGGAAGGGAGGGGAGGAGAGGCGCGGTAAGTGCATAATCATTTCATCAGTCCTT
AATAAACATATAGTGGGTGGGAGGGGCCGGCCGATATATGTAATCTGGCTAATAACTTGATCTG
GAGGGGCGGTGGTGGGGGAGCAGGGGAGGGGAGCCATCGCGGGGCGAGGCCGGTGGAGGGGAGG
TCGGCACCCTGATTGGCCGAGAGCGGGGAGTGGGCGGCGGTGAGGGGGCGGGGCCTTGTCGGAC
ACGCCCACCGCCCCCTCCCCACCACCGCACAGCGCTCTCCCGACTTTGAAATCTTAGTTAGAAG
CTGTCAGCCGTTGACAGAAGCTCTCAATCGGATGTAGTGAGGCGAGGCCGGCCAACATGCCACC
CAAGCGCAAAACGCTTAACGCTGCGGCTGAGGCTAACGCTCATGCCGACGGACACGCCGACGGA
AACGCCGACGGACACGTGGCCAATACGGCCGCGTCCTCGAATAATGCGAGGTTCGCTGATCTCA
CTAACATCGATACTCCGGGTCTGGGACCCACAACTACGACCCTGCTCGTGGAACCAGCACGCTC
AAAGCGTCAACGAGTGTCCCGCGCATGCGACCAGTGCCGTGCAGCCCGAGAGAAATGCGACGGA

ATACAGCCTGCGTGTTTCCCGTGCGTTTCCCAGGGAAGGTCCTGCACTTATCAGGCTTCGCCGA
AAAAGAGGGGAGTTCAAACCGGTTATATTCGTACGCTGGAGCTCGCCCTCGCCTGGATGTTTGA
AAATGTCGCGCGTTCCGAAGATGCCTTGCATAACCTCCTCGTCCGTGACGCCGGACAAGGATCA
GCTCTGCTCGTTGGTAAAGATTCGCCGGCTGCCGAGCGACTCCATGCCCGTTGGGCTACTAGCC
GTGTCAATAAGAGCATTACCCGCCTCCTCCGTCAGTTGGAGCTCCCTCCTACCGCCACGGCTAC
GGCCTCGATAATGCCGCACGTGATGGAGCAGCCTCTCAGTACCAGCATTAACCCCGTCAACGAC
CGCTTCAACGGTATTCCCAACCCCACTCCGTATAACTCCGATGCAGCTCTCGATGCTATCACTC
AGACCAACGATTATGGAAGCGTAAATACACATGGTATCCTCTCTACTTACCCGCCACCGGCTAC
GCACCTTAATGAAGCTTCCGTCGCTCTCGCTCCCGGTGGCGCCCCCCCCGACCGCCTCCTCCG
TATGTTGACAGCACGACCAATCACCCGCCGTACCACTCGAATCTGGTTCCAATGGCGAACTTTG
GTTACTCGACCGTTGATTACGATGCCATGGTTGACGATTTGGCTAGCATTGAATACACGGACGC
TGTGGATGTCGACCCACAGTTTATGACCAATCTGGGATTCGTTCCTGGATGTAACTTCTCCGAC
ATTAATACATACGAACAGTGATGACGTCTATCGATACCGTCGACTAAAGCCAAATAGAAAATTA
TTCAGTTCCTGGCTTAAGTTTTTAAAAGTGATATTATTTATTTGGTTGTAACCAACCAAAAGAA
TGTAAATAACTAATACATAATTATGTTAGTTTTAAGTTAGCAACAAATTGATTTTAGCTATATT
AGCTACTTGGTTAATAAATAGAATATATTTATTTAAAGATAATTGCGTTTTTATTGTCAGGGAG
TGAGTTTGCTTAAAAACTCGTTTAGATCCCGGTCACGTGTCACTGGGTCAGTGGGTAATCGCTT
ATCCTCGGATAAACAATTATCCTCACGGGTAATCGCTTATCCGCTCGGGTAATCGCTTATCCTC
GGGTAATCGCTTATCCTTGGGTAATCGCTTATCCTCGGATAAACAATTATCCTCACGGGTAATC
GCTTATCCGCTCGGGTAATCGCTTATCCTCGGGTAATCGCTTATCCTTGGGTAATCGCTTATCC
TCGGATAAACAATTATCCTCACGGGTAATCGCTTATCCGCTCGGGTAATCGCTTATCCTCGGGT
AATCGCTTATCCTTTCACGTTGGGACTCAGTGAGGAGGACCTGAATTCCTGCAGCCCGAGCGGA
GACTCTAGCGAGCGCCGGAGTATAAATAGAGGCGCTTCGTCTACGGAGCGACAATTCAATTCAA
ACAAGCAAAGTGAACACGTCGCTAAGCGAAAGCTAAGCAAATAAACAAGCGCAGCTGAACAAGC
TAAACAATCTGCAGTAAAGTGCAAGTTAAAGTGAATCAATTAAAAGTAACCAGCAACCAAGTAA

ATCAACTGCAACTACTGAAATCTGCCAAGAAGTAATTATTGAATACAAGAAGAGAACTCTGAATAGGGAATTGGGAATTCGTTAACAGATCGGGGGATCAATTCGTTAACATGGATAGCACCGAGGCAGTGATCAAGGAGTTCATGCGGTTCAAGGTGCACATGGAGGGCTCCATGAACGGCCACGAGTTCGAGATCGAGGGCGAGGGCGAGGGCCGCCCCTACGAGGGCACCCAGACCGCCAAGCTGAGGGTGACCAAGGGTGGCCCCCTGCCCTTCTCCTGGGACATCCTGTCCCCTCAGTTCATGTACGGCTCCAGGGCCTTCACGAAGCACCCCGCCGACATCCCCGACTACTGGAAGCAGTCCTTCCCCGAGGGCTTCAAGTGGGAGCGCGTGATGAACTTCGAGGACGGCGGCGCTGTGTCCGTGGCCCAGGACACCTCCCTGGAGGACGGCACCCTGATCTACAAGGTGAAGCTCCGCGGCACCAACTTCCCTCCTGACGGCCCCGTAATGCAGAAGAAGACAATGGGCTGGGAAGCATCCACCGAGCGGTTGTACCCCGAGGACGTCGTGCTGAAGGGCGACATTAAGATGGCCCTGCGCCTGAAGGACGGCGGTCGCTACCTGGCGGACTTCAAGACCACCTACAGGGCCAAGAAGCCCGTGCAGATGCCCGGCGCTTTCAACATCGACCGCAAGTTGGACATCACATCCCACAACGAGGACTACACCGTGGTGGAACAGTACGAACGCTCCGTGGCCCGCCACTCCACCGGCTAAAGCGGCCGCGACTCTAGATCATAATCAGCCATACCACATTTGTAGAGGTTTTACTTGCTTTAAAAAACCTCCCACACCTCCCCCTGAACCTGAAACATAAAATGAATGCAATTGTTGTTGTTAACTTGTTTATTGCAGCTTATAATGGTTACAAATAAAGCAATAGCATCACAAATTTCACAAATAAAGCATTTTTTTCACTGCATTCTAGTTGTGGTTTGTCCAAACTCATCAATGTATCTTAAGGTTCTGTGAATTCAGATCTCCGGGGATCTAATTCAATTAGAGACTAATTCAATTAGAGCTAATTCAATTAGGATTAATTCAATTAGAGACTAATTCAATTAGAGCTAATTCAATTAGGATTAATTCAATTAGAGACTAATTCAATTAGAGCTAATTCAATTAGGATCCAAGCTTATCGATTTCGAACCCTCGACCGCCGGAGTATAAATAGAGGCGCTTCGTCTACGGAGCGACAATTCAATTCAAACAAGCAAAGTGAACACGTCGCTAAGCGAAAGCTAAGCAAATAAACAAGCGCAGCTGAACAAGCTAAACAATCGAAGAATCAAAATGGTGAGCAAGGGCGAGGAGCTGTTCACCGGGGTGGTGCCCATCCTGGTCGAGCTGGACGGCGACGTAAACGGCCACAAGTTCAGCGTGTCCGGCGAGGGCGAGGGCGATGCCACCTACGGCAAGCTGACCCTGAAGTTCATCTGCACCACCGGCAAGCTGCCCGTGCCCTGGCCCACCCTCGTGACCACCCTGACCTACGGCGTGCAGTGCTTCAGCCGCTACCCCGACCACATGAA

GCAGCACGACTTCTTCAAGTCCGCCATGCCCGAAGGCTACGTCCAGGAGCGCACCATCTTCTTC
AAGGACGACGGCAACTACAAGACCCGCGCCGAGGTGAAGTTCGAGGGCGACACCCTGGTGAACC
GCATCGAGCTGAAGGGCATCGACTTCAAGGAGGACGGCAACATCCTGGGGCACAAGCTGGAGTA
CAACTACAACAGCCACAACGTCTATATCATGGCCGACAAGCAGAAGAACGGCATCAAGGTGAAC
TTCAAGATCCGCCACAACATCGAGGACGGCAGCGTGCAGCTCGCCGACCACTACCAGCAGAACA
CCCCCATCGGCGACGGCCCCGTGCTGCTGCCCGACAACCACTACCTGAGCACCCAGTCCGCCCT
GAGCAAAGACCCCAACGAGAAGCGCGATCACATGGTCCTGCTGGAGTTCGTGACCGCCGCCGGG
ATCACTCTCGGCATGGACGAGCTGTACAAGTAAAACTTGTTTATTGCAGCTTATAATGGTTACA
AATAAAGCAATAGCATCACAAATTTCACAAATAAAGCATTTTTTTCACTGCATTCTAGTTGTGG
TTTGTCCAAACTCATCAATGTATCTTGGTACCGGGCCCCCCCTCGAGGTCGACGGTATCGATAA
GCTTGATATCTATAACAAGAAAATATATATATAATAAGTTATCACGTAAGTAGAACATGAAATA
ACAATATAATTATCGTATGAGTTAAATCTTAAAAGTCACGTAAAAGATAATCATGCGTCATTTT
GACTCACGCGGTCGTTATAGTTCAAAATCAGTGACACTTACCGCATTGACAAGCACGCCTCACG
GGAGCTCCAAGCGGCGACTGAGATGTCCTAAATGCACAGCGACGGATTCGCGCTATTTAGAAAG
AGAGAGCAATATTTCAAGAATGCATGCGTCAATTTTACGCAGACTATCTTTCTAGGGTTAA

*piggybac* ITR: 238 bp(L), 305 bp(R)

*Oncopeltus invected 2.4-kb* upstream: 2435 bp

QF2: 1050 bp

*hsp70* terminator: 257 bp

15xQUAS: 290 bp

*Drosophila hsp70* promoter: 241 bp

mScarlet3: 672 bp

SV40 polyadenylation signal: 122 bp

Pax-6 homeodomain binding site: 11 bp x 9

*Drosophila hsp70* promoter (minimal): 125 bp

EGFP: 717 bp

SV40 polyadenylation signal: 121 bp

**Data S2. transposase protein sequence_261026**

>0xNLS-hyPBase

MGSTMPRSPEFAATMGSSLDDEHILSALLQSDDELVGEDSDSEVSDHVSEDDVQSDTEE
AFIDEVHEVQPTSSGSEILDEQNVIEQPGSSLASNRILTLPQRTIRGKNKHCWSTSKPT
RRSRVSALNIVRSQRGPTRMCRNIYDPLLCFKLFFTDEIISEIVKWTNAEISLKRRESM
TSATFRDTNEDEIYAFFGILVMTAVRKDNHMSTDDLFDRSLSMVYVSVMSRDRFDFLIR
CLRMDDKSIRPTLRENDVFTPVRKIWDLFIHQCIQNYTPGAHLTIDEQLLGFRGRCPFR
VYIPNKPSKYGIKILMMCDSGTKYMINGMPYLGRGTQTNGVPLGEYYVKELSKPVHGSC
RNITCDNWFTSIPLAKNLLQEPYKLTIVGTVRSNKREIPEVLKNSRSRPVGTSMFCFDG
PLTLVSYKPKPAKMVYLLSSCDEDASINESTGKPQMVMYYNQTKGGVDTLDQMCSVMTC
SRKTNRWPMALLYGMINIACINSFIIYSHNVSSKGEKVQSRKKFMRNLYMGLTSSFMRK
RLEAPTLKRYLRDNISNILPKEVPGTSDDSTEEPVMKKRTYCTYCPSKIRRKASASCKK
CKKVICREHNIDMCQSCF

EPVMKKRTYCTYCPSKIRRKASASCKKCKKVI = predicted endogenous NLS

>1xNLS-hyPBase

MGSTMPKKKRKVPRSPEFAATMGSSLDDEHILSALLQSDDELVGEDSDSEVSDHVSEDD
VQSDTEEAFIDEVHEVQPTSSGSEILDEQNVIEQPGSSLASNRILTLPQRTIRGKNKHC
WSTSKPTRRSRVSALNIVRSQRGPTRMCRNIYDPLLCFKLFFTDEIISEIVKWTNAEIS
LKRRESMTSATFRDTNEDEIYAFFGILVMTAVRKDNHMSTDDLFDRSLSMVYVSVMSRD
RFDFLIRCLRMDDKSIRPTLRENDVFTPVRKIWDLFIHQCIQNYTPGAHLTIDEQLLGF
RGRCPFRVYIPNKPSKYGIKILMMCDSGTKYMINGMPYLGRGTQTNGVPLGEYYVKELS
KPVHGSCRNITCDNWFTSIPLAKNLLQEPYKLTIVGTVRSNKREIPEVLKNSRSRPVGT
SMFCFDGPLTLVSYKPKPAKMVYLLSSCDEDASINESTGKPQMVMYYNQTKGGVDTLDQ
MCSVMTCSRKTNRWPMALLYGMINIACINSFIIYSHNVSSKGEKVQSRKKFMRNLYMGL
TSSFMRKRLEAPTLKRYLRDNISNILPKEVPGTSDDSTEEPVMKKRTYCTYCPSKIRRK
ASASCKKCKKVICREHNIDMCQSCF

PKKKRKV = SV40 NLS

EPVMKKRTYCTYCPSKIRRKASASCKKCKKVI = predicted endogenous NLS

```
>3xNLS-hyPBase
MGSTMPKKKRKVPKKKRKVPKKKRKVPRSPEFAATMGSSLDDEHILSALLQSDDELVGE
DSDSEVSDHVSEDDVQSDTEEAFIDEVHEVQPTSSGSEILDEQNVIEQPGSSLASNRIL
TLPQRTIRGKNKHCWSTSKPTRRSRVSALNIVRSQRGPTRMCRNIYDPLLCFKLFFTDE
IISEIVKWTNAEISLKRRESMTSATFRDTNEDEIYAFFGILVMTAVRKDNHMSTDDLFD
RSLSMVYVSVMSRDRFDFLIRCLRMDDKSIRPTLRENDVFTPVRKIWDLFIHQCIQNYT
PGAHLTIDEQLLGFRGRCPFRVYIPNKPSKYGIKILMMCDSGTKYMINGMPYLGRGTQT
NGVPLGEYYVKELSKPVHGSCRNITCDNWFTSIPLAKNLLQEPYKLTIVGTVRSNKREI
PEVLKNSRSRPVGTSMFCFDGPLTLVSYKPKPAKMVYLLSSCDEDASINESTGKPQMVM
YYNQTKGGVDTLDQMCSVMTCSRKTNRWPMALLYGMINIACINSFIIYSHNVSSKGEKV
QSRKKFMRNLYMGLTSSFMRKRLEAPTLKRYLRDNISNILPKEVPGTSDDSTEEPVMKK
RTYCTYCPSKIRRKASASCKKCKKVICREHNIDMCQSCF
```

PKKKRKVPKKKRKVPKKKRKV = 3x SV40 NLS

EPVMKKRTYCTYCPSKIRRKASASCKKCKKVI = predicted endogenous NLS

>5xNLS-hyPBase

MGSTMPKKKRKVPKKKRKVPKKKRKVPKKKRKVPKKKRKVPRSPEFAATMGSSLDDEHI
LSALLQSDDELVGEDSDSEVSDHVSEDDVQSDTEEAFIDEVHEVQPTSSGSEILDEQNV
IEQPGSSLASNRILTLPQRTIRGKNKHCWSTSKPTRRSRVSALNIVRSQRGPTRMCRNI
YDPLLCFKLFFTDEIISEIVKWTNAEISLKRRESMTSATFRDTNEDEIYAFFGILVMTA
VRKDNHMSTDDLFDRSLSMVYVSVMSRDRFDFLIRCLRMDDKSIRPTLRENDVFTPVRK
IWDLFIHQCIQNYTPGAHLTIDEQLLGFRGRCPFRVYIPNKPSKYGIKILMMCDSGTKY
MINGMPYLGRGTQTNGVPLGEYYVKELSKPVHGSCRNITCDNWFTSIPLAKNLLQEPYK
LTIVGTVRSNKREIPEVLKNSRSRPVGTSMFCFDGPLTLVSYKPKPAKMVYLLSSCDED
ASINESTGKPQMVMYYNQTKGGVDTLDQMCSVMTCSRKTNRWPMALLYGMINIACINSF
IIYSHNVSSKGEKVQSRKKFMRNLYMGLTSSFMRKRLEAPTLKRYLRDNISNILPKEVP
GTSDDSTEEPVMKKRTYCTYCPSKIRRKASASCKKCKKVICREHNIDMCQSCF

PKKKRKVPKKKRKVPKKKRKVPKKKRKVPKKKRKV = 5x SV40 NLS

EPVMKKRTYCTYCPSKIRRKASASCKKCKKVI = predicted endogenous NLS

#### Supplementary Video Legends

**Supplemental Video 1.** Ventral view of a developing *O. fasciatus* embryo showing ubiquitous expression of H2B:mScarlet driven by the endogenous actin promoter identified herein. Over this 36 hour time course, katatrepsis (second inversion of the embryonic axes such that the head of the embryo moves from the posterior of the egg (right) to the anterior of the egg (left) (Panfilio, 2008)), and appendage elongation occur. The Green Fire Blue look up table implemented in Fiji (Schindelin et al., 2012) shows the highest intensity pixels in white, intermediate intensity pixels in green, and the lowest intensity pixels in black (see Figure 5D). Download Link

**Supplemental Video 2.** Dorsal view of a developing *O. fasciatus* embryo showing ubiquitous expression of H2B:mScarlet driven by the endogenous actin promoter identified herein. Over this 36 hour time course, the end of anatrepsis (first inversion of the embryonic anteroposterior axis such that the head of the embryo moves from the anterior of the egg (right) to the posterior of the egg (left)), and the whole of katatrepsis (second inversion of both anteroposterior and dorsoventral embryonic axes relative to those of the egg) occur (Panfilio, 2008). The Green Fire Blue look up table implemented in Fiji (Schindelin et al., 2012) shows the highest intensity pixels in white, intermediate intensity pixels in green, and the lowest intensity pixels in black (see Figure 5D). Download Link

**Supplemental Video 3.** Lateral view of a developing *O. fasciatus* embryo showing ubiquitous expression of H2B:mScarlet driven by the endogenous actin promoter identified herein. Over this 36 hour time course, the end of anatrepsis and the whole of katatrepsis (Panfilio, 2008) occur as in Supplemental Video 3. From this lateral viewpoint, emerging muscular contractions of the embryo are apparent beginning at approximately 30 hours, the elongating appendages on the ventral side of the embryo (bottom), and the dorsal movement (towards the top) of the lateral ectoderm that is required for dorsal closure, are evident. The Green Fire Blue look up table implemented in Fiji (Schindelin et al., 2012) shows the highest intensity pixels in white, intermediate intensity pixels in green, and the lowest intensity pixels in black (see Figure 5D).

Download Link
